# Phosphoproteomic dysregulation drives tumor proliferation in Cushing’s disease

**DOI:** 10.1101/2025.09.28.679056

**Authors:** David T. Asuzu, Dhruval Bhatt, Dustin Mullaney, Debjani Mandal, Diana Nwokoye, Sheelu Varghese, Daniela Tortoza Lopez, Nikhil Ramavenkat, Kory Johnson, Abdel Elkahloun, Zied Abdullaev, Kenneth Aldape, Dragan Maric, Clarisse Quignon, Nasir S. Malik, Joseph P. Steiner, Yan Li, Susan Wray, Christina Tatsi, Lynnette K Nieman, Prashant Chittiboina

## Abstract

Pituitary adenomas constitute up to 20% of primary brain tumors, yet somatic mutations are only found in 15% of pituitary adenomas. Epigenomic dysregulation has been proposed as a tumorigenic mechanism in pituitary adenomas causing Cushing’s disease (CD). We created paired datasets of human CD adenomas and en-route margin adult human pituitary glands and assayed their chromatin accessibility, DNA methylation, transcriptomic, proteomic and phospho-proteomic landscapes. In CD adenomas, we found epigenetic reactivation of a neurodevelopmental phosphoprotein program typically lost in the post-natal pituitary gland. CD cells overexpressed *PPP1R17*, a potent endogenous inhibitor of the ubiquitous protein phosphatase PP2A. Mechanistically, *PPP1R17* overexpression in normal murine pituitary cells recapitulated the adenoma phenotype, and PPP1R17-mediated tumorigenesis was reversible using an FDA-approved small molecule PP2A agonist both in-vitro and in-vivo. Our findings highlight aberrant peptide phosphorylation as a targetable mechanism in CD.

**Significance statement:** Cushing’s disease (CD) causes significant morbidity and mortality despite best medical and surgical treatment. Surgery is the mainstay of treatment, but carries perioperative risks and is frequently followed by remission. There is a paucity of effective medical treatments, due in part to a limited understanding of tumor mechanisms. The majority of CD adenomas are wild-type, with no known causal mutations. Our study identifies phosphoproteomic dysregulation as a mechanism of CD tumorigenesis common to wild-type and mutant CD adenomas. We target this pathway in-vivo and in-vitro using an FDA-approved small molecule. Our study proposes a novel therapeutic strategy for patients with CD.

## Introduction

Eukaryotic cellular homeostasis is tightly governed by the balance of protein kinases and phosphatases, which control phosphorylation levels and enzymatic activity of key intracellular proteins ^1,2^. Disruption of this kinase-phosphatase balance has been reported underlying malignant tumors driven by discrete mutations ^3,4^. In non-malignant tumors, kinase dysregulation has been described as a mechanism of neoplastic tumor progression following the acquisition of mutations ^5^. Little is known about early aberrant phosphorylation events in non-malignant tumor formation, or about epigenetic mechanisms underlying such dysregulation.

The PP2A family of serine/threonine phosphatases plays a critical role in the regulation of signal transduction cascades, cell cycle regulation, cell morphology and development ^6^. PP2A is regulated by the relative abundance of its regulatory subunits, which control its substrate specificity, subcellular localization and catalytic activity ^7–10^. The catalytic subunit (PP2Ac) is further regulated by methylation or phosphorylation. Several endogenous PP2A inhibitors have been identified, which either bind directly to PP2Ac or target specific PP2A holoenzymes and prevent dephosphorylation of PP2A substrates ^11^. PPP1R17 is an endogenous PP2A inhibitor both in its phospho-PPP1R17 and dephospho-PPP1R17 forms ^8^, however, its role in pituitary tumorigenesis has not been previously investigated.

The pituitary gland is the site of sporadic adenomas (benign tumors) in up to 35% of the general population ^12,13^. Despite its small size (∼500mg), adenomas of the pituitary gland comprise 20% of all primary brain tumors ^14^. Even after successful removal, histologically benign adenomas can recur in up to 30% of patients, causing long-term morbidity ^15^. Cushing’s disease (CD) is caused by pituitary adenomas that hypersecrete adrenocorticotropin (ACTH). CD adenomas are often small (2 - 10 mm) at diagnosis, minimally distort the pituitary architecture and have a predictable effect on the hormonal axes (biochemical testing confirms the presence of and remission from CD) ^16–18^. Non-surgical treatments are limited, therefore, understanding mechanisms of tumorigenesis may improve our ability to target these tumors ^19^.

A minority of CD adenomas (total ∼40%) have somatic mutations in *USP8* (∼30%), *USP48* or *BRAF* ^20–22^. Epigenetic dysregulation has been hypothesized to contribute to CD tumorigenesis via promoter hypomethylation of *POMC*, *ESR1* and *CASP8* ^23,24^, via downregulation of antiproliferative micro-RNAs miR-15a, miR-16 and miR-26a ^25,26^, and via increased glucocorticoid resistance triggered by loss of *HDAC2* and Swi/Snf chromatin remodeling complex subunit *Brg1* ^27^. However, the chromatin accessibility landscape of CD remains unknown. Previously, we applied advances in surgical sampling and in-vitro modeling to identify anti-apoptotic mechanisms in wildtype CD adenomas ^28^. Here, we investigated common epigenetic mechanisms underlying CD adenomas using a multi-omics approach.

## Results

### PPP1R17 is overexpressed in CD adenomas

We performed single nucleus (sn) multiome analysis on nuclei extracted from surgically annotated (**Supplemental video**) freshly dissociated CD adenomas versus *en-route,* adjacent cells in the tumor margin (total 37,263 nuclei; **Figure 1A, Supplemental Table S1**). We included wildtype and *USP8*-mutated CD adenomas in this study (**Figure 1B, Supplemental Table S1**) and used marker-based cell-type classification of single nuclei ^29,30^ that corresponded well with unsupervised clustering ^31,32^. Using a validated set of expression markers ^33^, we identified canonical pituitary cell classes in each sample and generated snRNAseq gene signatures for corticotrophs, lactotrophs, somatotrophs, endothelial cells, gonadotrophs, leukocytes and folliculostellate cells (FSCs; **Figure 1C, Supplemental Figure S1, Supplemental Table S2**). Importantly, we distinguished between core CD corticotrophs versus adjacent corticotrophs in the tumor margin (**Figure 1C**). We found marked similarities between cell-type specific gene signatures derived from snRNAseq compared to single cell RNAseq ^34^ ^28^. In the integrated snRNAseq dataset, most cells within the tumor core were corticotrophs, whereas corticotrophs comprised only a small fraction of cells in the tumor margin (**Figure 1D**, left two panels).

**Figure 1.**
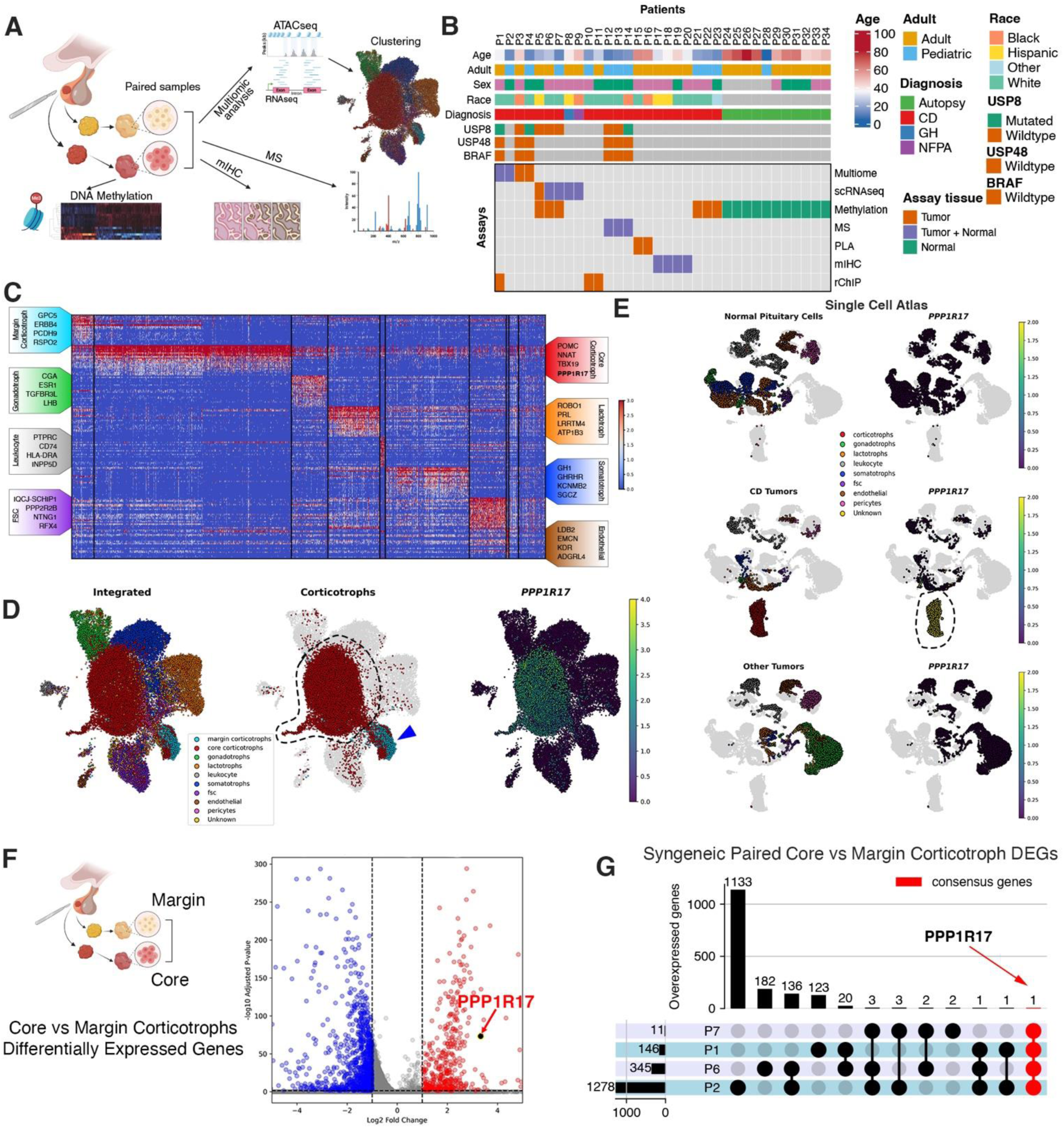
*PPP1R17* is overexpressed in CD adenomas. **A.** Study schema. Pituitary adenomas and en-route adjacent margin tissues (when available) were surgically annotated and processed separately for single-nucleus multiome (ATACseq plus RNAseq) analysis, Mass spectrometry (MS), multiplex immunocytochemistry (mIHC) and DNA methylation. **B.** Summary of human surgical samples and investigations included in the current study. **C.** Marker gene expression in the integrated multiome dataset and their corresponding cell classes (see also: Supplemental Figure S1 and Supplemental Table S2). **D.** Left panel: UMAP projections of distinct nuclei from snRNAseq analysis using patient-derived samples including post-natal pituitary cells in the CD tumor core versus margin. Nuclei were clustered according to their cell-type identity. Middle panel: CD core specimens are comprised mostly of corticotrophs. Right panel: PPP1R17 expression is largely restricted to core CD corticotrophs. **E.** scRNAseq analysis 28 evaluating PPP1R17 expression (right column) mapped over pituitary cell types (left column). Top row: post-natal pituitary cells from the tumor margin. Middle row: CD core adenomas. Bottom row: non-CD adenomas (1 growth hormone and 1 non-functioning pituitary adenoma). **F.** Volcano plot from snRNAseq data showing differentially expressed genes between CD core versus margin corticotrophs. **G.** UpSet plot with pairwise comparisons between syngeneic CD core versus margin corticotrophs from the same patients evaluated by snRNAseq (P1 versus P2) or scRNAseq (P6 versus P7). CD = Cushing’s disease.

Next, we evaluated for genes differentially overexpressed in CD core versus margin cells using snRNAseq. *PPP1R17* was predominantly expressed in CD core corticotrophs (**Figure 1D**, right panel), a finding which was observed both in our previously published scRNAseq dataset ^28^ as well as in the current snRNAseq dataset (**Figure 1E-F**). Using a combination of trajectory inference and RNA velocity using CellRank ^35^, the fate probability transition from margin to tumor phenotype was strongly linked to *PPP1R17* overexpression (**Supplemental Figure S2).** Furthermore, when we limited our analysis to syngeneic tumor-margin samples from the same patients (P1 and P2 snRNAseq, and P6 and P7 scRNAseq), *PPP1R17* was the only gene differentially overexpressed in CD adenomas versus syngeneic margin pituitary glands (**Figure 1G; Supplemental Table S4**)

### Mechanisms of PPP1R17 overexpression in CD adenomas

To better understand the mechanisms underlying *PPP1R17* overexpression, we leveraged our snATACseq data and mapped the corticotroph-restricted fragment coverage (binned to 1 Mb) over the entire genome. We found increased chromatin accessibility in adenoma corticotrophs compared to margin corticotrophs (**Figure 2A**). We then mapped fragments to ‘peaks’ specific to corticotrophs, and individual genes. Upon differential analysis, we found increased coverage at peak regions, regions proximal to transcription start sites and distal to transcription end sites in CD adenoma corticotrophs (**Figure 2B**).

**Figure 2:**
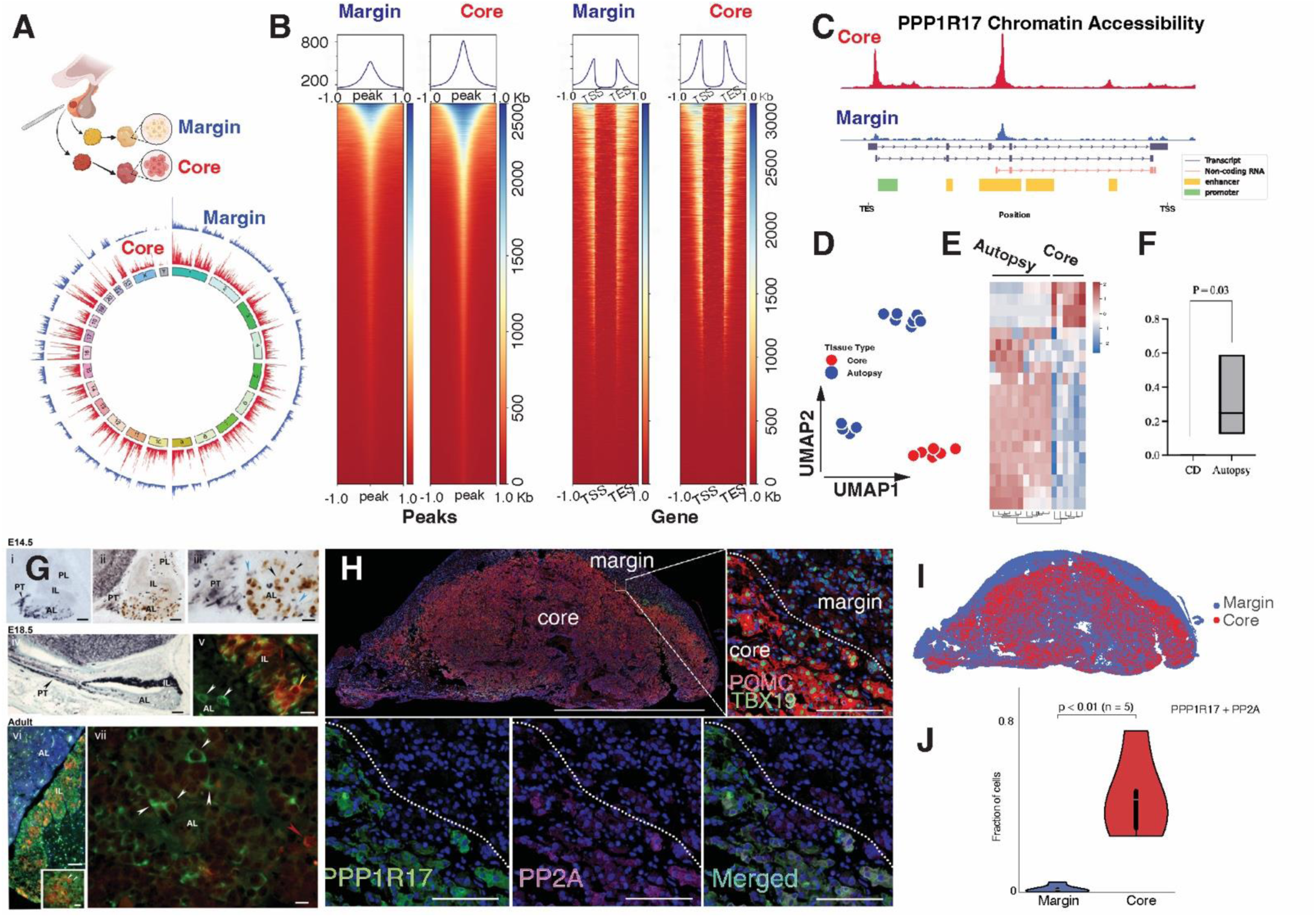
Altered global and *PPP1R17* locus-specific chromatin accessibility in CD adenoma cells. **A.** Fragment coverage over the entire genome in the single-nucleus multiome dataset. Fragment coverage of corticotroph cells is averaged over 4 syngeneic adenoma core-margin pairs and two additional adenomas, binned to 1mBase, and mapped at the same scale for adenoma corticotrophs (red) and adjacent margin corticotrophs (blue). **B.** Waterfall plots mapping reads per kilobase million (RPKM)-normalized fragment coverage in corticotrophs over peak (left), and gene (right) regions. In each panel, the right column is the adenoma corticotroph map and the left is for corticotrophs derived from the adjacent margin. **C.** Chromatin accessibility at the *PPP1R17* locus in adenoma corticotrophs (red track) versus corticotrophs derived from the adjacent margin (blue track), plotted at the same scale. Chromatin accessibility data were derived from 2 syngeneic adenoma margin pairs and two additional CD adenomas and averaged across compartments. **D-E.** Differentially methylated regions in CD adenomas (n = 6) versus autopsy-derived normal pituitary glands (n = 11). **F.** Bisulfite conversion followed by methylation-specific PCR evaluating methylation at the *PPP1R17* promoter in CD adenomas (n = 5) versus autopsy-derived normal pituitary glands (n = 5, subset from E). **G.** i: Alpha GSU staining in mouse E14.5 embryos highlights pars tuberalis (PT) and anterior lobe (AL). ii – iii: Pomc staining (brown) is mostly restricted to the AL, and ppp1R17 staining (blue)is found in the PT and AL (blue arrowheads). Iv: By E18.5, ppp1R17 staining (blue) is mostly restricted to the intermediate lobe (IL) and PT, with only few select AL cells retaining ppp1R17 positivity. V: Pomc (green) staining AL and IL, and ppp1R17 (red) restricted to the IL. Vi-vii: minimal Ppp1R17 expression in the adult mouse (red) in the AL, with residual staining in the IL. Scale bar = 50µM. **H.** Representative mIHC image (P17) of a whole-mounted CD adenoma and its adjacent margin tissues (i-ii) corticotroph-specific POMC and TBX19 expression. PPP1R17 (iii), and its canonical substrate PP2Ac (iv) co-expression (v) assessed within the identified adenoma. Scale bar = 50µM. **I.** Quantitative analysis of mIHC signal intensity across 18 channels (except PPP1R17 and PP2Ac) identifies core/adenoma cells distinct from margin/normal cells. **J.** Averaged proportional co-expression of PPP1R17 and PP2Ac in margin/normal (1.68%) and core/adenoma cells (45%) across 4 samples.

We identified 1,094 differentially accessible peaks (606 up in the tumor core, 488 up in the adjacent margin) and 517 differentially accessible genes (150 up in the tumor core, 367 up in the adjacent margin; **Supplemental Figure S3A, Supplemental Table S3**). Comparative gene-set enrichment (GSEA) analysis of gene expression and chromatin accessibility profiles revealed a significant overlap in pathways (n = 67 GO pathways) including cytoplasmic translation and oxidative phosphorylation (**Supplemental Figure S3B-C, Supplemental Table S5**). Several transcription factor motifs were differentially accessible in CD adenomas, including NF-Y and SP1 (**Supplemental Table S6**). NF-Y is involved in chromatin remodeling, promotes cell proliferation and declines during differentiation ^36,37^. SP1 is an essential embryonic transcription factor responsible for regulating cell cycle, methylation-free CpG islands, and formation of active chromatin structures ^38^. Correspondingly, several NF-Y and SP1 targets were differentially overexpressed in adenoma corticotrophs (**Supplemental Figure S3D-E, Supplemental Table S7**).

We focused on the *PPP1R17* locus and identified increased chromatin accessibility at the *PPP1R17* promoter (**Figure 2C**). We also found increased chromatin accessibility at an internal enhancer from which a novel lncRNA *PPP1R17-203* is transcribed (**Supplemental Figure S4**). *PPP1R17-203* shares sequence similarity with *MAP7* which plays a role in several cancer types ^39^, and BLASTN revealed interactions between *PPP1R17-203* and several other lncRNAs (**Supplemental Table S8**). The role of *PPP1R17-203* in CD tumorigenesis remains to be explored.

Pituitary adenomas can be clustered according to their DNA methylation profiles ^23^. We asked if promoter DNA hypomethylation contributed to *PPP1R17* overexpression. DNA methylation profiling on tissues from 6 CD adenomas versus 11 autopsy-derived pituitary adenomas revealed widespread DNA methylation changes (**Figure 2D-E, Supplemental Table S9**). Bisulfite conversion and methylation-specific PCR from a subset of DNA methylation samples including 5 CD adenomas versus 5 autopsy-derived human pituitaries identified DNA hypomethylation at the *PPP1R17* promoter (P = 0.03; **Figure 2F**). Analysis of TCGA data similarly found *PPP1R17* promoter hypomethylation in several tumor types including hepatocellular carcinoma and head and neck squamous cancer (**Supplemental Figure S5A)**. Importantly, there was no concurrent over- or under-expression of other PP1 or PP2A subunit genes (**Supplemental Figure S5B**). These data suggest that epigenetic dysregulation contributes to *PPP1R17* overexpression in CD adenomas.

### Identification of upstream drivers of PPP1R17 overexpression in CD

*PPP1R17* is expressed in neural progenitor cells during human fetal development but is lost in adulthood except for cerebellar Purkinje cells and occasional cells in the hypothalamus and pons/medulla ^40,41^. We found robust Ppp1r17 expression in E14.5 mouse embryonic pituitary glands, which decreased markedly by E18.5 (**Figure 2G iv - v**) and diminished further in the adult mouse anterior pituitary lobe where we found only sparse and low expression (**Figure 2G vi - vii**). In contrast, PPP1R17 protein was abundant within the CD adenoma compartment and colocalized with its target protein PP2Ac, a ubiquitous serine-threonine phosphatase ^42^ and tumor suppressor ^43^ (**Figure 2H**, **Supplemental Figure S6**). Upon quantitative analysis (Ward method^44^), a significant (p<0.01) proportion (45%) of core/adenoma cells co-expressed PPP1R17 and PP2Ac compared to margin/normal cells (**Figure 2I**) across all samples tested (**Figure 2J, Supplemental Figure S7**).

We next assessed proteins associated with the accessible *PPP1R17* promoter in CD adenomas by reverse-ChIP (rChIP) using CRISPR dCas9-3XFLAG and a gRNA specific to the *PPP1R17* promoter to pull down promoter-associated proteins for Mass spectrometry ^45^ (**Supplemental Figure S8**). Control experiments excluded the *PPP1R17* targeting gRNA. Proteins bound to the *PPP1R17* promoter in CD adenomas (**Supplemental Table S10)** involved mRNA splicing and processing pathways (**Supplemental Figure S8**). Several histone variants including H1.2, H1.4, H2A.2, H3.2 were also enriched at the *PPP1R17* promoter, as well as HP1BP3 which plays a key role in heterochromatin organization. Additionally, several members of the RAS oncogene family were enriched at the *PPP1R17* promoter, including RAB10, RAB15, RAB1A, RAB2B and RAB3B. Our snRNAseq data verified *RAB3B* overexpression in CD corticotrophs (**Supplemental Table S2**). Taken together, these findings suggest that increased RAS signaling ^46^ and chromatin remodeling events contribute to *PPP1R17* reactivation in CD corticotrophs

### PPP1R17 interacts with PP2A in CD adenomas

PPP1R17 has been identified as an endogenous inhibitor of protein phosphatase 1 (PP1), and of tumor-suppressor protein phosphatase 2A (PP2A).^47,48^ GSEA of differentially expressed genes in tumor corticotrophs (compared to normal corticotrophs) revealed a shift towards PP2A (LINCS L1000 KO Consensus Sigs geneset^49^; **Supplemental Figure S9A**) and it’s inhibitor okadaic acid (LINCS L1000 Chem Pert Consensus Sigs geneset^50^; **Supplemental Figure S9B**) compared to PP1 and it’s inhibitor calyculin. Likewise, applying a robust geneset (Reactome pathway 2024 geneset) revealed signal only from the PP2A complex in tumor corticotrophs (**Supplemental Figure S9C**), suggesting biologically relevant PP2A inhibition was restricted to tumor corticotrophs. The most common mode of PP2A inhibition in human tumors is via overexpression of other known endogenous PP2A inhibitors including CIP2A, PME1 and SET ^51,52^. Other than *PPP1R17*, we did not find transcriptional overexpression of other endogenous PP2A inhibitors in adenoma corticotrophs (**Supplemental Figure S10**). Known endogenous inhibitors PME1 and SET interact with PP2Ac at Arg89 and Tyr127. AlphaFold Multimer ^53^ predicted binding of human PPP1R17 to the PP2Ac similarly (**Figure 3A**). We verified PPP1R17 colocalization with PP2Ac in human CD adenomas by proximity ligation assay (**Figure 3B**) in addition to mIHC (**Figure 2H**). Based on the expression of B regulatory genes including *PPP12R2A-D* (**Supplemental Figure S5B**), we suspect that the predominant PP2A holoenzyme in human corticotrophs is B55.

**Figure 3:**
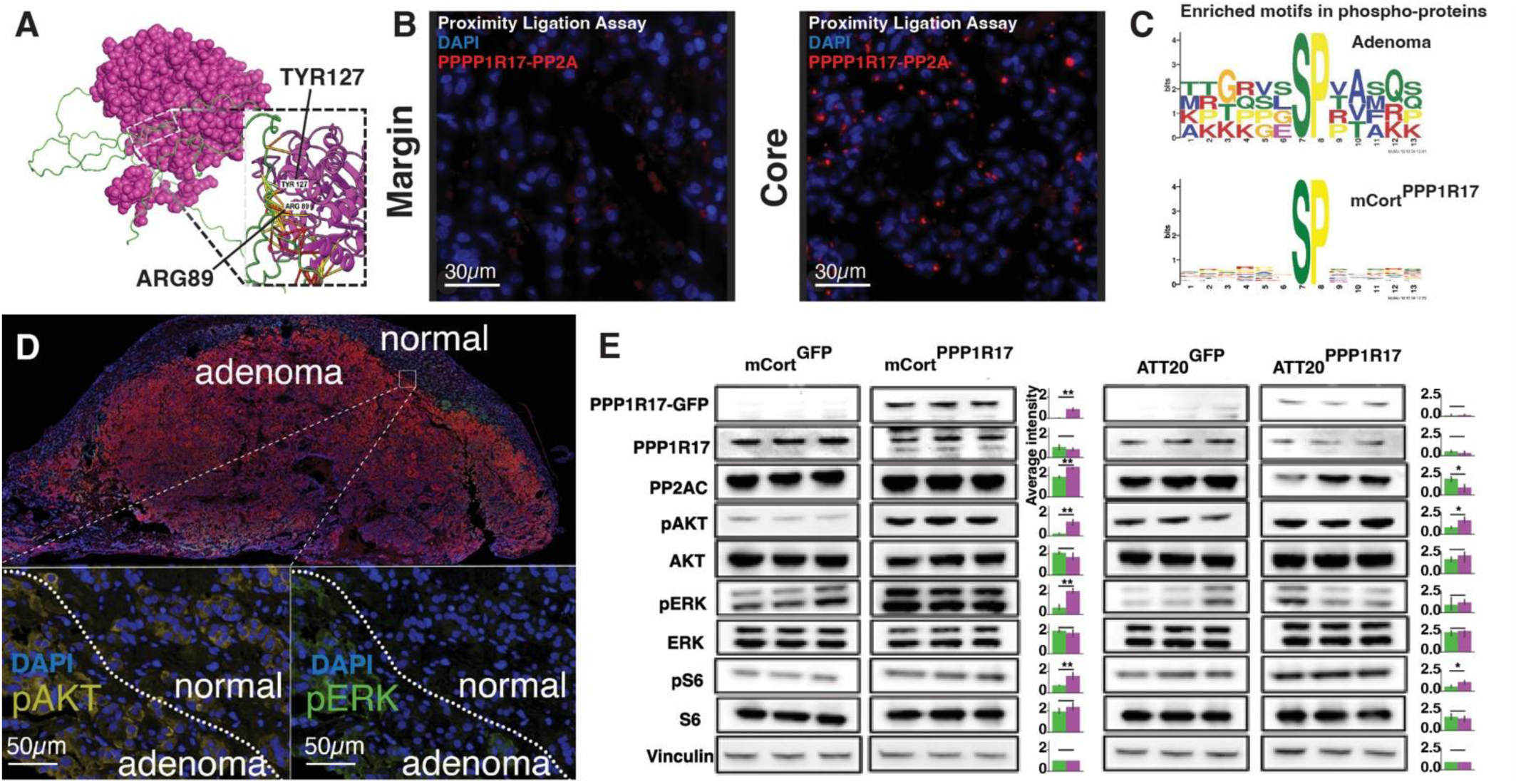
PPP1R17 inhibits PP2A phosphatase activity in CD. **A.** AlphaFold Multimer modeling predicting binding of human PPP1R17 to PP2Ac subunit. **B.** Representative proximity ligation assay (PLA) in margin (left) versus CD adenoma core (right) using antibodies for PP2Ac and PPP1R17. Red puncta indicate peptides in proximity. **C.** MoMo motif analysis of preferentially hyperphosphorylated and dephosphorylated motifs in CD adenoma core versus margin (n = 3/group; top panel) or in mCort^PPP1R17^ versus mCort^GFP^ cells (n = 5/group; bottom panel). **D.** Representative multiplexed immunohistochemistry staining of a CD adenoma (P17) and its adjacent margin. Inset identifies the region of interest that is magnified to demonstrate pERK and pAKT at the margin of core and margin. **E.** Western blots of mCort cells and ATT-20 cells transiently transfected with lentiviral plasmids encoding *PPP1R17* versus *GFP* evaluating phosphorylation of PP2A targets AKT, ERK and S6. (n = 3/condition; bar graphs with t-tests; * p < 0.05; ** p < 0.01; blank – not significant).

### PPP1R17 reactivation induces tumorigenic signaling in adenomas

We further explored the downstream effects of PP2A inhibition by performing quantitative phosphoproteomic profiling where we labeled syngeneic, paired CD adenomas and their adjacent margin with Tandem Mass Tag (TMT) reagents, enriched phosphopeptides using TiO2 and iMac methods, and acquired LC-MS/MS data on peptides with and without phosphoenrichment. We normalized phosphopeptides to their corresponding peptides. We performed MoMo motif analysis in CD adenomas versus adjacent margin ^54^ and identified preferential hyperphosphorylation of peptides with motifs xxSPxx and DxxxSxx (**Figure 3C, top panel, Supplemental Figure S11A-B**). Correspondingly, several PP2A targets were hyperphosphorylated in CD adenomas including pERK and pAKT (**Figure 3D**). We identified hyperphosphorylation of peptides involved in RNA splicing and chromosome condensation (**Supplemental Table S11**).

We next validated the activation of tumorigenic pathways in-vitro using mouse anterior pituitary cells (mCort) to serve as baseline normal cells. We stably overexpressed *PPP1R17* tagged to GFP in mCort cells (mCort^PPP1R17^; **Supplemental Figure S12**). With *PPP1R17* over-expression, there was no significant up or down regulation of PP1 and PP2A peptides. Following PPP1R17 pulldown, we found enrichment of phosphatase subunits including Ppp1r2c (11-fold, p = 2.17e-13), and Ppp2r2d (1.5-fold, p = 0.001) (**Supplemental Figure S13**).

Motif analysis in mCort^PPP1R17^ versus mCort^GFP^ cells also identified preferential hyperphosphorylation of peptides with the xxSPxx motif (**Figure 3C, bottom panel; Supplemental Figure S11C-D**). mCort^PPP1R17^ cells exhibited enrichment of phosphoproteins similarly associated with mitotic spindles, RNA splicing and G2-M checkpoint regulation (**Supplemental Table S12**). We interrogated phosphoproteomic modulation of known tumorigenic pathways with western blotting. mCort cells transiently transfected using *PPP1R17* lentiviral plasmids showed elevated pAKT (p = 0.002), pERK (p = 0.001) and pS6 ( p = 0.009) compared to mCort cells transfected with *GFP* (**Figure 3E, left panel**). In ATT-20 cells, a model for CD ^55^, transient transfection using *PPP1R17* lentiviral plasmids similarly resulted in elevated pAKT (p = 0.04) and pS6 (p = 0.02) compared to *GFP* controls (**Figure 3E, right panel**).

### PPP1R17 overexpression promotes a transcriptional upregulation program in CD

In addition to its role as a cytoplasmic phosphatase, PP2A can dephosphorylate nuclear RNA polymerase II via the INTAC complex and globally attenuate transcription ^56^. We therefore hypothesized that chronic *PPP1R17* overexpression activates RNA PolII, resulting in increased transcription and differential protein abundance in CD adenomas. mCort cells stably overexpressing PPP1R17 (mCort^PPP1R17^) showed increased pRNA PolII (S2, S5, S7) and increased total RNA PolII compared to mCort^GFP^ cells (**Figure 4A**) suggesting transcriptional derepression. Bulk RNAseq analysis of mCort^PPP1R17^ compared to mCort^GFP^ cells revealed upregulation of genes associated with MYC activation, unfolded protein response and epithelial-mesenchymal transition (**Figure 4B, Supplemental Table S13**), pathways previously associated with CD adenomas ^19,28,57^. We observed significant overlap between genes overexpressed in human CD adenomas and mCort ^PPP1R17^ cells, especially in genes involved in protein translation and peptide biosynthesis (**Supplemental Table S14**).

**Figure 4:**
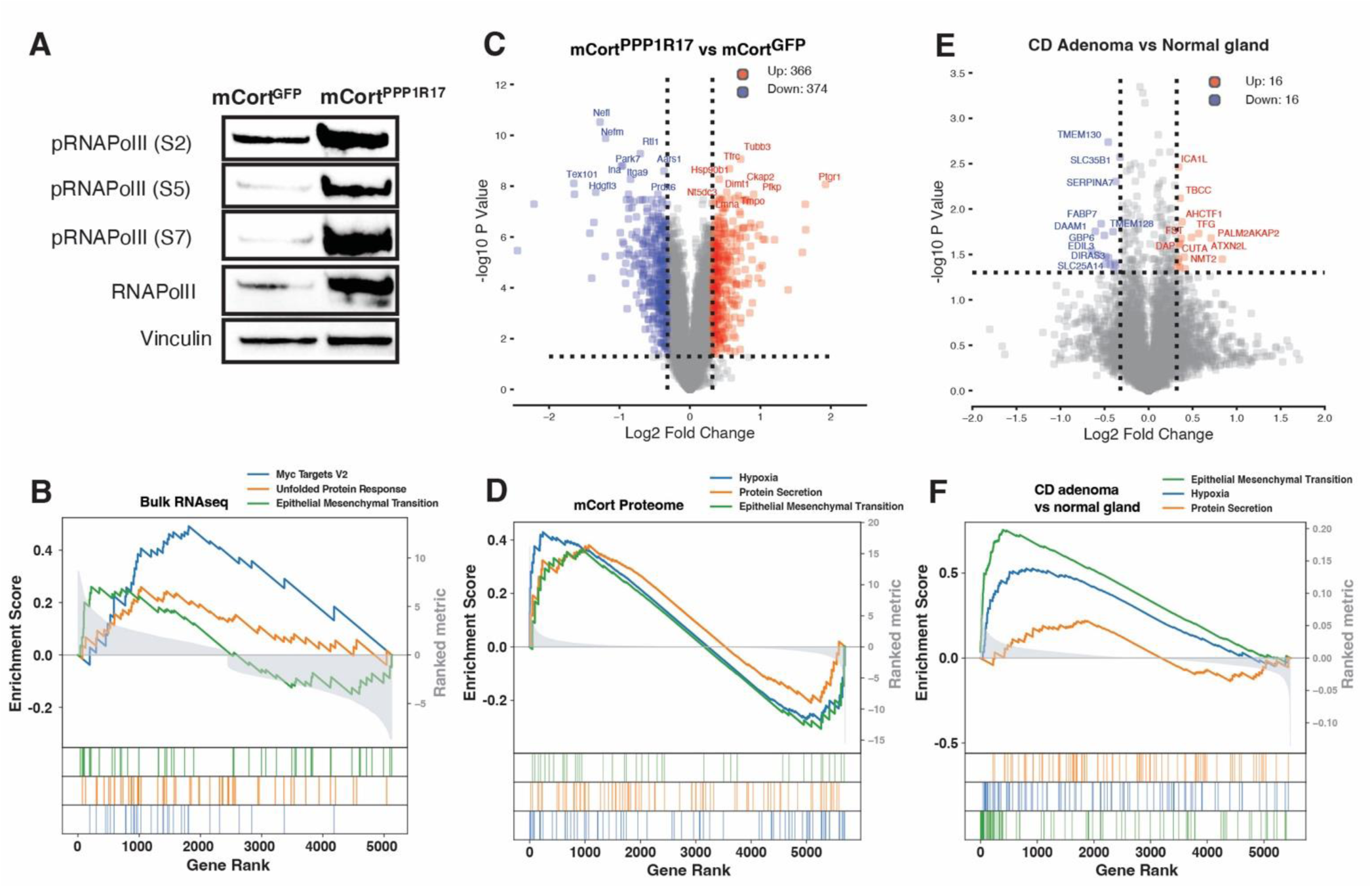
*PPP1R17* overexpression in-vitro recapitulates tumorigenic programs. **A.** Representative Western blots evaluating phosphorylation of RNA PolII (S2, S5 and S7) as well as total RNA PolII in mCort^PPP1R17^ versus mcort^GFP^ cells. **B.** Geneset enrichment (GSEA) analysis of bulk RNAseq comparison of mCort^PPP1R17^ (left) versus mCort^GFP^ cells (n = 3/group) showing differential enrichment of several pathways. **C.** Proteomic analysis of mCort^PPP1R17^ versus mCort^GFP^ cells (n = 5/group). **D.** GSEA analysis of peptides differentially abundant in mCort^PPP1R17^ (left) versus mCort^GFP^ cells (n = 5/group) highlighting enrichment of pathways for hypoxia, protein secretion and epithelial-mesenchymal transition. **E.** Proteomic analysis of CD adenomas versus their adjacent margin (n = 3/group). **F.** GSEA analysis of peptides differentially abundant in CD adenomas (left) versus adjacent margin (n = 3/group) highlighting enrichment of pathways for epithelial-mesenchymal transition, hypoxia and protein secretion.

Proteomic analysis of mCort^PPP1R17^ compared to mCort^GFP^ cells (**Figure 5C**) identified upregulation of peptides involved in epithelial-mesenchymal transition, hypoxia and protein secretion (**Figure 4C-D, Supplemental Table S15**). Accordingly, we found increased EGFR staining in CD adenomas (**Supplemental Figure S14**). Proteomic analysis of CD adenomas versus their adjacent margin tissues similarly revealed enrichment of proteins involved with epithelial-mesenchymal transition, hypoxia and protein secretion (**Figure 4E-F, Supplemental Table S16**), suggesting that *PPP1R17* overexpression potentiates a hormone secreting tumor program.

**Figure 5:**
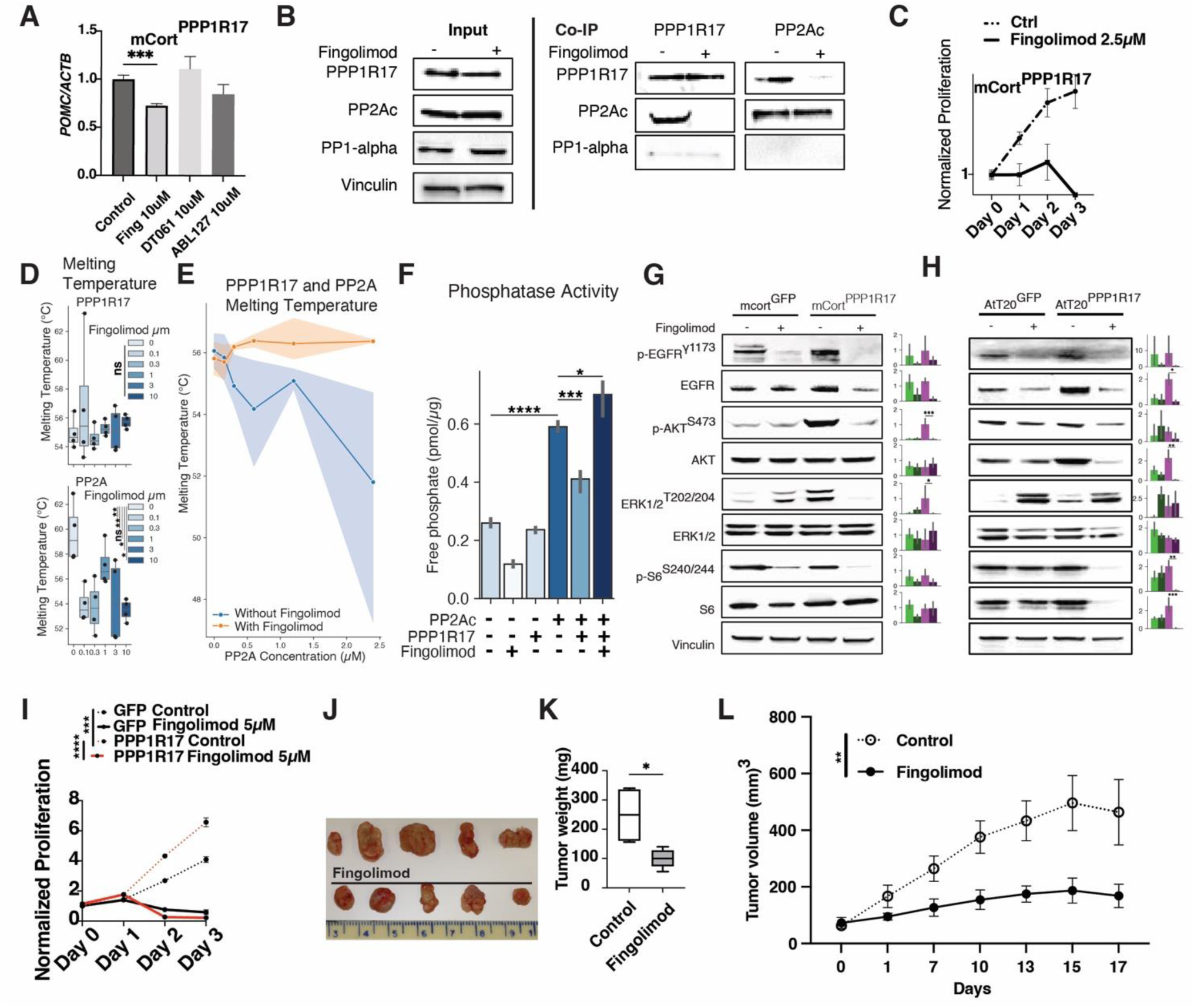
PPP1R17 upregulation is a targetable mechanism in CD. **A.** qRT-PCR of POMC normalized to ACTB in mCort^PPP1R17^ treated with fingolimod, DT061 or ABL127 (10 µM each; n = 3 technical replicates, representative image from n = 3 biological replicates; *** P < 0.05). **B.** Co-immunoprecipitation in At-T20 cells using anti-PPP1R17 and anti-PP2A antibodies, with and without fingolimod (5 µM, 2 hours). Western blots show PPP1R17, PP2Ac, and PP1-alpha (n = 3 replicates). **C.** Proliferation of mCort^PPP1R17^ vs. mCort^GFP^ cells after treatment with fingolimod (2.5 µM, n = 8/group). **D.** Melting temperatures of purified recombinant human PPP1R17 and PP2Ac in the presence and absence of fingolimod (0.1-10 uM), determined by Thermofluor assay (n = 4 replicates per condition; box-and-whisker plots). **E.** Melting temperature of PPP1R17 in the presence of increasing PP2Ac concentrations (0–2.5 µM), with or without fingolimod (3 µM; n = 4 replicates per condition). Shaded regions represent 95% confidence intervals. **F.** Western immunoblotting for phosphorylation of EGFR and PP2A targets (AKT, ERK and S6) in mCort^PPP1R17^ cells versus mCort^GFP^ cells with and without fingolimod treatment (5 µM for 24 hours; representative of 3 biological replicates). **G.** Western immunoblotting of phosphorylation status of PP2A targets EGFR and AKT in mCort^PPP1R17^ versus mCort^GFP^ cells with and without fingolimod treatment (5 µM for 24 hours; representative of 3 biological replicates). Quantitation of 3 biological replicates represented with bar plot (ANOVA; Post-hoc Tukey test; * p = 0.01, ** p = 0.001, *** p = 0.0004). **H.** Western immunoblotting for phosphorylation of EGFR and PP2A targets (AKT, ERK, and S6) in AtT-20PPP1R17 versus AtT-20GFP cells with and without fingolimod treatment (2.5 µM for 24 hours; representative of 3 biological replicates). Quantitation of 3 biological replicates represented with bar plot (ANOVA; Post-hoc Tukey test; * p = 0.04, ** p = 0.001, *** p = 0.0004). **I.** Proliferation of AtT-20PPP1R17 versus AtT-20GFP cells after treatment with fingolimod (5 µM) or DMSO control (n = 8/group; *** P < 0.001, **** P < 0.0001). **J.** Representative photograph of AtT-20 xenograft tumors from athymic NU/NU mice treated with fingolimod (1 mg/kg i.p.) or normal saline for 17 days. **K.** Tumor weights following fingolimod or saline treatment (n = 5/group; * P < 0.05). L. Tumor volumes over 17 days in fingolimod-versus saline-treated mice (n = 5/group; ** P < 0.05).

### PPP1R17 overexpression is a targetable mechanism of adenoma pathogenesis

PP2A agonists interrupt the interaction between PP2A and its endogenous inhibitors, a strategy that may be of clinical benefit ^58–60^. However, this strategy has not been explored in the context of hormone-secreting adenomas. We tested PP2A agonists DT061, ABL127, and fingolimod in mCort^PPP1R17^ cells and found that fingolimod suppressed *POMC* transcription (**Figure 5A**). Using anti-PPP1R17 and anti-PP2Ac (catalytic subunit for the PPP2A complex) for co-immunoprecipitation, we confirmed the interaction of PPP1R17-PP2Ac in At-T20 cells (**Figure 5B**). Additionally, we found evidence of minimal PP1 interaction with PPP1R17 in At-T20. Next, we confirmed that fingolimod treatment prevented PPP1R17 binding to PP2Ac, but not to PP1 (**Figure 5B**) in AtT-20 cells. Fingolimod inhibited cell proliferation in mCort^PPP1R17^ cells (**Figure 5C**) but not in mCort^GFP^ cells (**Supplemental Figure S15A**).

We purified recombinant human PPP1R17 and PP2Ac and performed Thermofluor protein thermal shift assays to confirm the mechanism of fingolimod action. We titrated purified PPP1R17 and obtained reliable melting temperature measurements at 1μg of protein. Fingolimod treatment decreased the melting temperature of PP2A (> 2.5 C; p < 0.01 at 0.1 µM fingolimod) but not PPP1R17 (**Figure 5D**). PP2Ac resulted in a concentration-dependent decrease in the PPP1R17 melting temperature, with a half-maximal effect between 1.5 and 2.5 μg of PP2A (**Figure 5E**), verifying direct interaction between PPP1R17 and PP2Ac. This thermal shift was attenuated by addition of fingolimod, a competitive PP2A agonist (**Figure 5E**). Purified, recombinant PP2Ac retained significant serine/threonine phosphatase activity in cell free lysates as measured by free phosphate abundance (p < 0.0001). PPP1R17 attenuated PP2Ac phosphatase activity (p = 0.002), an effect that was rescued by fingolimod (10 µM, 30 min; p = 0.01) (**Figure 5F**).

Phosphoproteomic analysis showed that treatment of mCort^PPP1R17^ cells with fingolimod selectively decreased phosphorylation of peptides with the xxSPxx motif (**Supplemental Figure S15B).** Dephosphorylated peptides were involved in cell cycle regulation and RNA polymerase II transcription (**Supplemental Figure S15C; Supplemental Table S17**). We next interrogated the specific peptides that fingolimod reversed the phosphorylation following PPP1R17 overexpression (hypo-hyper-hypo phosphorylation pattern). Fingolimod reversed PPP1R17-mediated hyperphosphorylation in several proteins (**Supplemental Figure S15D**) involved in MYC signaling and G2-M checkpoint regulation (**Supplemental Figure S15E**, **Supplemental Table S17**). We confirmed that fingolimod treatment reversed AKT (pAKTS473; p = 0.0004) and ERK (pERK1/2T202/2024; p = 0.004) hyperphosphorylation in mCort^PP1R17^ cells (**Figure 5F**) and induced apoptotic markers Bid, Bbc3 and Hrk (Supplemental **Figure 15F**).

As in mCort cells, stable PPP1R17 overexpression in AtT-20 cells induced tumorigenic signaling (**Figure 5H**) and proliferation compared to AtT-20^GFP^ cells (**Figure 5I**), and both were reversed by 5μM fingolimod treatment**. Specifically, we found a statistically significant reduction in phosphorylated AKT (p = 0.001).** Since parental AtT-20 cells have a tumor-like phenotype, we found that fingolimod also reduced the viability of AtT-20^GFP^ cells. We generated AtT-20 xenografts in athymic NU/NU mice which resulted in adrenal hypertrophy (**Supplemental Figure S18A**). Fingolimod treatment (1mg/Kg) led to a significant decrease in tumor size (**Figure 5J**), tumor weights (100 vs 249 mg, P = 0.05; **Figure 5K**) and measured volumes (final volume 168 vs 463 mm^3^, P = 0.05; **Figure 5L; Supplemental Figure S15B**). There were no significant changes in skin thickness or body weights in tumor-bearing mice treated with fingolimod versus saline control for 14 days (**Supplemental Figure S15C**). Taken together, these findings identify PP2A agonism as a therapeutic strategy for CD adenomas.

## Discussion

### PPP1R17 is an endogenous PP2A inhibitor in pituitary adenomas causing Cushing’s disease

Our findings demonstrate that epigenetic dysregulation of the kinase-phosphatase balance can underlie human tumors. In this study, we identified epigenetic reactivation of *PPP1R17,* an endogenous inhibitor of the protein phosphatase PP2A, as a driver of CD tumorigenesis. PPP1R17 activity was associated with widespread dysregulation of the phosphoproteome. Reversal of this phenotype using the PP2A agonist fingolimod rescued cellular hyperproliferation and induced apoptosis in a tumor model. Kinase-phosphatase imbalance due to somatic gene amplification of *PRKAR1A* has been implicated in adrenal tumors causing Cushing’s syndrome ^61^. Here, we find convergent epigenetic dysregulation along a different pathway leading to kinase-phosphatase imbalance in pituitary adenomas causing Cushing’s disease.

Epigenetic modulation of gene expression is a tumorigenic mechanism whereby environmental cues impinge upon the transcriptome without requiring changes in the DNA coding sequence ^62^. The majority of CD adenomas are not associated with canonical mutations of known targets *BRAF*, *USP8* or *USP48* ^22,63^, and studies on CD epigenomics are lacking. Our multiomic approach combining single-nucleus RNA and ATACseq with DNA methylation profiling identified epigenetic *PPP1R17* reactivation in CD adenomas, highlighting a role for gene-environment interactions in CD. Our reverse-ChIP results suggested histone subunit recruitment and active chromatin remodeling at the *PPP1R17* promoter, which agrees with other reports on histone partitioning as a mechanism of tumor progression ^64^. Further studies are needed to determine precisely how environmental factors modify histone recruitment in CD.

### PPP1R17 reactivation as a targetable mechanism of CD

PPP1R17, also known as G-substrate due to its preferential phosphorylation by cGMP-dependent protein kinase, is found almost exclusively in human cerebellar Purkinje cells where it plays a role in the induction of long-term depression ^7,65^. In the rat brain, Ppp1r17 is also restricted almost entirely to Purkinje cells of the cerebellum, although low levels are detectable in the paraventricular region of the hypothalamus and in the pons/medulla ^41^. Unlike in adults, *PPP1R17* is highly expressed in neural progenitor cells in the developing human cortex ^66–68^. This expression pattern is restricted to mammals and is not seen in ferrets or mice of comparable ages ^69^.

In the prenatal macaque and human cortex, *PPP1R17* is localized to the outer and inner subventricular zones, and colocalizes with markers of intermediate progenitor cells and dividing cells ^69^. RNA-seq time-course studies also confirmed that *PPP1R17* expression is restricted to neural progenitor cells during fetal development in humans ^40^, therefore its overexpression in CD likely reflects epigenetic reactivation through coordinated increase in chromatin accessibility and DNA hypomethylation. Other studies have recently described similar epigenetic reactivation of developmental genes in the pathogenesis of pulmonary hypertension ^70^, however this reactivation has not previously been described in pituitary tumors.

The proximate mechanism of PPP1R17 reactivation in CD adenomas is still not fully understood. Whereas our results suggest underlying epigenetic dysregulation via promoter hypomethylation, increased chromatin accessibility and RAS signaling, the underlying index event is still unclear. A response to PP2A upregulation in the adenoma is possible, although we did not identify transcriptional upregulation of other PP2A agonists *CIP2A*, *PME1* and *SET* in CD adenomas. It is notable that the pituitary gland resides outside the blood brain barrier adjacent to the median eminence with its leaky vasculature, which allows for prompt homeostatic responses to environmental stressors. This architecture also allows transmission of blood-borne pathogens and other environmental agents which trigger hormone responses from the hypothalamus and pituitary gland. Given that epigenomic changes frequently occur in response to environmental cues, we speculate that cumulative exposure to environmental insults or infectious agents could create an epigenomic milieu favoring tumorigenesis via *PPP1R17* dysregulation. Additional studies are needed to investigate this hypothesis.

### PPP1R17 overactivity induces widespread phosphoproteomic changes

PP2A is a member of the family of serine / threonine phosphoprotein phosphatases (PPPs), which unlike metal-dependent phosphatases (PPMs) or protein tyrosine phosphatases (PTPs), utilize conserved motifs for metal-dependent catalysis. PPPs include PP1, PP2A, PP2B, PP4, PP5, PP6 and PP7, and often consist of holoenzymes with regulatory subunits which control their location and specificity. While PP2A inactivation is linked to uncontrolled growth in several cancers^71, 71^ aberrant PP1 signaling also contributes to tumorigenesis^72^. Several other PPPs are upregulated in the tumor context, and associated with poor prognoses^73^, suggesting context-specific activation. Our data highlights the role of PP2A inhibition in CD, however a role for other protein phosphatases cannot be excluded in CD.

Overexpression of PP2A inhibitors has been identified as an oncogenic mechanism in multiple human cancers, including *SET* overexpression in CML, AML, T-cell ALL, B-cell CLL, non-small cell lung cancer and Wilms tumors ^51,74–76^, and *CIP2A* overexpression in hepatocellular, breast, colorectal, ovarian, cervical, prostate, lung, head and neck cancer ^52,77–79^. Small molecules that activate PP2A (SMAPs) are an emerging class of antitumor agents with therapeutic potential as combinatorial agents in lung cancer models ^80,81^, as well as in multiple myeloma, leukemia and colorectal cancer ^60^. Our results show the therapeutic potential of PP2A agonism using fingolimod in CD both in-vitro and in-vivo. Interestingly, *PPP1R17* overexpression in mouse cortical neurospheres inhibits the G1-S transition and results in decreased proliferation ^69^. In our study, *PPP1R17* overexpression in mature corticotrophs (mCort) similarly resulted in prolonged G0/G1 but led to accelerated S phase and increased proliferation in mCort^PPP1R17^ cells, indicating cellular context-specific effects.

In addition to its role in protein dephosphorylation, PP2A was recently shown to exert direct transcriptional control by dephosphorylating RNA PolII via an integrator complex ^56,60,82^. We noted phosphorylation of RNA PolII in mCort^PPP1R17^ cells, with resultant differential expression of several genes. Furthermore, we noted significant overlap between the gene expression profile of mCort^PPP1R17^ cells and CD adenomas, indicating that several transcriptomic changes in CD likely result from PPP1R17-mediated PP2A inhibition. Future studies will investigate whether PPP1R17 may be partly mediating kinase-phosphatase imbalance through its interactions with PRKACA, which plays a role in the formation of adrenal tumors ^83^.

Our study is limited by the small number of patient samples included, partly due to limited availability of *en-route* adult human pituitary tissues in the tumor margin. Fingolimod also possesses PP2A-independent effects, including its effects on immune cells ^84^, which may limit its therapeutic window for patients with CD. Future studies will develop and test more specific PPP1R17 inhibitors for use in humans.

## Conclusions

Our study highlights the role of kinase-phosphatase disequilibrium in the pathogenesis of benign tumors, specifically in adrenocorticotropin secreting pituitary adenomas that cause CD. We identified epigenetic reactivation of *PPP1R17*, an endogenous inhibitor of PP2A, as a central driver of CD tumorigenesis irrespective of the somatic mutation status. The therapeutic potential of targeting this imbalance is underscored by the efficacy of PP2A agonism in reversing the tumorigenic effects of *PPP1R17* overexpression in-vitro and in-vivo. This work advances our understanding of pituitary adenoma formation and suggests a promising avenue for therapeutic intervention in benign tumors.

## Supplemental Figure Legends

**Supplemental Figure S1.** Related to Figure 1D. UMAPs of canonical cell types in each individual tumor sample.

**Supplemental Figure S2.** Tumor corticotroph lineage tracing analysis. A. CellRank and scVelo analysis of patient sample P2. i. scVelo velocities projected onto UMAP embedding of all cells in both tumor and normal compartments colored by cell type. ii. scVelo velocities projected onto UMAP embedding of tumor corticotrophs, normal corticotrophs, and folliculostellate cells (FSC). iii. UMAP of corticotrophs from tumor and normal compartments. Cells are colored by CellRank fate probabilities, which is the likelihood that a given cell will transition toward the terminal population. iv. CellRank identified initial cell states. v. CellRank identified terminal cell states. vi. Transition matrix showing the fate probabilities of each macrostate. vii. Relative expression of relevant genes in each macrostate plotted over latent time. B. CellRank and scVelo analysis of patient sample P1. scVelo velocities projected onto UMAP embedding of tumor corticotrophs, normal corticotrophs, and FSCs (left), scVelo velocities projected onto UMAP embedding of tumor and normal corticotrophs (center), scVelo velocities projected onto UMAP embedding of tumor and normal corticotrophs with cells colored by relative *PPP1R17* expression (center). C. CellRank and scVelo analysis of patient sample P2. scVelo velocities projected onto UMAP embedding of tumor and normal corticotrophs. Cells colored by relative *PPP1R17* expression. D. CellRank and scVelo analysis of patient sample P3. scVelo velocities projected onto UMAP embedding of tumor corticotrophs and FSCs (left), scVelo velocities projected onto UMAP embedding of only corticotrophs (center), scVelo velocities projected onto UMAP embedding of corticotrophs with cells colored by relative *PPP1R17* expression (center). E. CellRank and scVelo analysis of patient sample P3. scVelo velocities projected onto UMAP embedding of tumor corticotrophs and FSCs (left), scVelo velocities projected onto UMAP embedding of only corticotrophs (center), scVelo velocities projected onto UMAP embedding of corticotrophs with cells colored by relative *PPP1R17* expression (center).

**Supplemental Figure S3.** Related to Figures 2A and 2B. **A.** Multiome analysis comparing snRNAseq and snATACseq in CD corticotrophs versus adjacent normal corticotrophs. **B.** GSEA analysis of pathways upregulated by snRNAseq in CD corticotrophs versus adjacent normal corticotrophs. **C.** Left panel: overlap between genes differentially expressed (GEX) and genes differentially accessible (ATAC) in CD corticotrophs. Right panel: GSEA analysis of pathways differentially accessible by snATACseq in CD corticotrophs. **D.** Comparison of chromatin accessibility (left panels) and gene expression (right panels) for NF-Y target gene CALM1 in CD corticotrophs (red, tumor) versus adjacent normal corticotrophs (blue, normal). **E.** Differential chromatin accessibility (left panels) and gene expression (right panels) for SP1 target gene SCIN in CD corticotrophs (red, tumor) versus adjacent normal corticotrophs (blue, normal).

**Supplemental Figure S4.** The novel lncRNA PPP1R17-203 is transcribed from an internal *PPP1R17* site. A. Long-range interactions predicted for PPP1R17-203. B. Predicted 3D structure of PPP1R17-203. Source: https://bioinformaticslab.erc.monash.edu/linc2function

**Supplemental Figure S5.** A. DNA methylation analysis from the TCGA database demonstrating *PPP1R17* promoter hypomethylation in multiple solid tumors including hepatocellular carcinoma and head and neck squamous cancer. B. Volcano plot and dotplot from snRNAseq data showing differentially expressed genes between CD core/adenoma versus margin/normal corticotrophs.

**Supplemental Figure S6.** PPP1R17 expression is confined to CD adenomas. Related to Figure 2H. A. Multiplex immunohistochemistry of CD adenoma and surrounding normal pituitary gland (i) showing PPP1R17 expression (green) compared to corticotroph-specific POMC expression (red). PP2Ac expression (purple) staining is compared to PPP1R17 staining within the adenoma (iii – v). Scale bar = 50µM. B. Representative H&E of normal pituitary gland and CD adenoma demonstrating PPP1R17 staining. CD = Cushing’s disease. H&E = hematoxylin and eosin.

**Supplementary Figure S7:** Co-expression and quantitation of PPP1R17 and PP2A in tissue samples. Related to Figures 2H, 2H and 2J. A. As in Figure 2I, expression based segmentation of multiplexed immunohistochemistry (mIHC) was performed on all available samples. Adenoma and normal cells were classified according to expression of 18 proteins (except PPP1R17 and PP2Ac). Red indicates core/adenoma cells, and blue indicates margin/normal cells. B. Barplots with statistical testing results (Fisher’s exact test; **** is p < 0.0001).

**Supplemental Figure S8.** Schematic for reverse-ChIP performed using CRISPR dCas9-3XFLAG-biotin and a gRNA targeting the *PPP1R17* promoter to pull down promoter-bound peptides for mass spectrometry. GSEA analysis for proteins bound to the *PPP1R17* promoter reveals enrichment for RNA splicing and RNA processing.

**Supplemental Figure S9.** Geneset enrichment analysis (GSEA) for corticotroph cells. Core/adenoma (Left in GSEA plot) are compared to corticotroph cells from margin/normal (Right in GSEA plot) compartments. A. LINCS L1000 knockdown consensus signature geneset, with plotting of transcriptional signature of knockdowns of catalytic subunits of PP2A (PPP2CA), and PP1 (PPP1CA). B. Transcriptional signatures of PP2A inhibition (okadaic acid), and PP1 (calyculin) plotted with the LINCS L1000 Chem Pert Consensus Sigs geneset. C. Reactome pathway 2024 geneset identified transcriptional signature of PP2A signaling.

**overexpressed in CD.** UMAP projections demonstrating expression patterns of *PPME1* and *SET* in *POMC*-overexpressing CD adenoma cells. AlphaFold multimer predicts PME1 and SET interaction with PP2Ac at Arg89 and share an additional interaction site at Tyr127 with PPP1R17. UMAP = uniform manifold approximation and projection. CD = Cushing’s disease.

**Supplemental Figure S11. A.** TMT LC-MS/MS quantification of phosphopeptide/peptide ratios in core CD adenomas versus margin (n = 3/group) shows several hyperphosphorylated and dephosphorylated targets. **B**. MoMo motif analysis of preferentially hyperphosphorylated and dephosphorylated motifs in CD adenoma core versus margin (n = 3/group). **C**. TMT LC-MS/MS quantification of phosphopeptide/peptide ratios in mCort^PPP1R17^ versus mCort^GFP^ cells (n = 5/group). **D.** MoMo motif analysis of preferentially hyperphosphorylated and dephosphorylated motifs in mCort^PPP1R17^ versus mCort^GFP^ cells (n = 5/group).

**Supplemental Figure S12. Characterization of murine corticotroph (mCort) cells transfected with lentiviral PPP1R17. A**. Lentiviral vector overexpressing *PPP1R17* tagged to *GFP*. Stable transfection in mCort cells was verified by qRT-PCR. **B.** Western blot characterizing PPP1R17 (26 kDa) and PPP1R17-GFP proteins (54kDa) in mCort cells. **C.** Stable *PPP1R17* overexpression in mCort cells (mCort^PPP1R17^) showed hyperproliferation compared to mCort^GFP^ cells (n = 8/group). mCort^PPP1R17^ cells also showed acceleration of the cell cycle.

**Supplemental Figure S13.** Liquid chromatography/mass spectroscopy (LC/MS/MS) for peptides following PPP1R17 over-expression in mCort cells. Detectable PP1 and PP2 complex peptides, and Ppp1r17 are highlighted in green. **A.** Volcano plots comparing peptide abundance in mCort^PPP1R17^ cells compared with mCort^GFP^ cells. PP1 and PP2A complex subunit peptides are highlighted in green. **B.** Volcano plot for enrichment of peptides belonging to PP1 (Ppp1r12c) and PP2A (Ppp2r2d) with Ppp1r17 pull-down followed by LC/MS/MS.

**Supplemental Figure S14.** EGFR is overexpressed in CD. Representative mIHC image of whole-mounted CD adenoma and adjacent normal gland (i) with corticotroph-specific POMC expression. (ii-iv) EGFR expression (green) was compared to POMC expression (red). Scale bar = 200 µM.

**Supplemental Figure S15. A.** Effect of fingolimod and DT061 on proliferation in mCort^GFP^ cells; n = 8/group. **B.** MoMo analysis showed differential dephosphorylation of peptides with the xxSPxx motif in mCort^PPP1R17^ cells after treatment with 48 hours of fingolimod 10μM (n = 5/group). **C.** GSEA analysis revealed dephosphorylation of peptides involved in cell cycle signaling and RNA Polymerase II transcription in mCort^PPP1R17^ cells after 48 hours of fingolimod 10μM. **C.** Heatmap of individual proteins hyperphosphorylated in mCort^PPP1R17^ but rescued by 48-hour fingolimod treatment. **E.** GSEA analysis of rescued phosphoproteins. F. Heatmap of apoptotic pathway marker expression from a custom PCR panel in mCort^PPP1R17^ cells treated with DMSO, fingolimod, DT061, or ABL127 (10 µM; n = 3 technical replicates). **B – D** are the results of TMT labeling and mass spectrometry; n = 5/group; one-way ANOVA P < 0.05.

**Supplemental Figure S16. A**. Implantation of ATT20 cells into the flanks of nude mice induced adrenal gland hypertrophy (n = 5/group), which did not return to baseline after fingolimod treatment (n = 4). **B.** Nude mice bearing ATT20 flank xenografts formed tumors after 10-12 days. After tumor establishment, intraperitoneal fingolimod 1mg/kg daily for 17 days led to decreased tumor size. **C.** Fingolimod treatment had no effect on skin thickness or animal body weight.

## Acknowledgements

This study was supported by the Intramural Research Programs of the National Institute of Neurological Disorders and Stroke, *Eunice Kennedy Shriver* National Institute for Child Health, Human Development, and the National Institute of Diabetes and Digestive Kidney Diseases and National Human Genome Research Institute, Bethesda, MD, USA.

## Author contributions

DTA study design, bioinformatics, data analysis and interpretation, paper drafting and editing. DB data analysis, bioinformatics, figure generation, paper editing..DMu data analysis, bioinformatics, figure generation, paper editing. DMan *in-vitro* experiments. DN *in-vitro* experiments. NR bioinformatics, data analysis. KJ bioinformatics. AE high throughput sequencing. ZA in vitro assays, data analysis. KA pathological assessment. DMar immunofluorescence. CQ immunofluorescence. SW immunofluorescence, data analysis and interpretation, paper editing. NKL protein assays. NSM in vitro studies. JPS protein analysis, data interpretation. YL proteome/phosphoproteome analysis, data interpretation. SW animal studies and immunostaining. LKN study design, data interpretation. CT study design, data interpretation. PC study design, data analysis and interpretation, figure generation, study supervision, and manuscript editing.

## Data and materials availability

### Lead contact

Further information and requests for resources and reagents should be directed to and will be fulfilled by the lead contact: Prashant Chittiboina, MD, MPH. Tenure Track Investigator, Neurosurgery Unit for Pituitary and Inheritable Diseases, National Institute of Neurological Diseases and Stroke, National Institutes of Health. 10 Center Drive, Room 3D20, Bethesda, MD 20892-1414. Phone: (301) 496-5728. Fax: (301) 402-0380.

### Materials availability

This study did not generate new unique reagents.

### Data and code availability

High throughput sequencing data generated from this study has been uploaded to Geo. Custom analysis toolkits and code developed for this study has been uploaded to Github. Additional methods are available upon request from the corresponding author.

## List of Supplemental Materials

Supplemental Methods

Supplemental Figures S1 through S16

Supplemental Tables S1 through S17

Supplemental video

## Materials and methods

### Resource Availability

#### Materials availability

This study did not generate new unique reagents.

### Experimental model and study participant details

#### Patients

Surgical samples were obtained from non-consecutive patients with a biochemical diagnosis of CD undergoing pituitary surgery by a single surgeon (PC) between 2006 and 2023 (**Supplemental Table S1**). This study was conducted at the National Institutes of Health (NIH) Clinical Center in Bethesda, MD and approved by the combined neuroscience Institutional Review Board of the National Institute of Neurological Disorders and Stroke, Bethesda, MD (clinicaltrials.gov identifier NCT00060541). Each patient gave written informed consent.

Hypercortisolism in CD patients was diagnosed as previously described ^85,86,87^. All patients underwent a sub-labial trans-sphenoidal approach for microscope-assisted selective adenectomy. Adenomas were identified pre-operatively using high-resolution magnetic resonance imaging (MRIs) with Pituitary protocol, and their locations were verified intraoperatively by visual inspection. Adenomas were removed by micro-surgical dissection along a circumferential pseudo-capsular plane. Pituitary adenoma was removed en-bloc within its pseudocapsule wherever possible ^88^. The en-route or adjacent non-viable adjacent normal pituitary gland was removed separately and annotated for analysis. Research specimens were labeled separately and fresh-frozen for further study.

#### Animal studies

All procedures were approved by National Institute of Neurological Disorders and Stroke, Animal Care and Use Committee and performed in accordance with National Institutes of Health guidelines. Adult pregnant NIH Swiss female mice were euthanized with CO2 at two gestation ages, embryonic day (E) 14.5 and E18.5. Embryos and pituitaries of the adult dams were collected, fixed in 4% formaldehyde (Sigma)/0.1M phosphate buffer saline (PBS, 24h), transferred to 30% sucrose (48hr, 4°C), embedded in OCT (Sakura Tissue-Tek) and stored at - 80°C until sectioning. Sections (pituitary, 12µm; E14.5 and E18.5, 16µm) were cut on a cryostat (Leica Biosystem CM 3050S).

For in-vivo tumor injections, 4-5 week-old female athymic nude mice (NU/NU) were purchased from Charles River Laboratories and housed under standard conditions in a pathogen-free facility with ad libitum access to food and water. Mice were grouped into untreated mice, mice injected with cells, and mice injected with cells and treated with fingolimod. Number of mice per group were determined based on power calculations assuming effect size of 30% based on in-vitro data. Cell suspensions of 7 x 10^6^ cells in 100mL PBS were injected into the flank subcutaneously using 25G needles.

After allowing 10-12 days for tumor establishment, tumor-bearing mice were randomized into DMSO versus fingolimod treatment groups. Fingolimod was dissolved in normal saline and injected intraperitoneally at a daily concentration of 1mg/Kg body weight. Mice were monitored daily for clinical signs of distress by dedicated animal handlers at the NIH Animal Research Facility in keeping with NIH IACUC guidelines. A licensed veterinarian from the NIH was on call 24 hours for any animal emergencies. At the end of the treatment period, surviving mice were euthanized humanely using carbon dioxide according to AVMA Guidelines for the Euthanasia of Animals and NIH IACUC guidelines.

#### Corticotroph cells and constructs

Corticotroph cells (mCort) were harvested as previously described ^28^ from female BALB/c mice aged 6-8 weeks (Taconic Biosciences, USA) ^33^ Lentiviral vectors encoding *PPP1R17-GFP* or *GFP*-only were obtained from Origene (Cat# RC207007L4V and PS100093). 10 IFUs/5×10^6^ cells were applied in normal media to mCORT or AtT-20 cells for 48 hours, followed by antibiotic selection for 2 weeks using 1mM Puromycin (Thermo Fisher #A1113802). GFP positivity was confirmed in surviving cells, and *PPP1R17* expression was verified using qRT-PCR and Western immunoblotting.

#### Single Nucleus Multiome analysis

CD adenomas and adjacent normal pituitary glands were embedded in OCT and stored at -80°C. Nuclei were isolated from the frozen tissues using an adaptation of a previously established protocol ^89^. Nuclei were then immediately processed using a Single Cell Multiome ATAC + Gene Expression Assay (10X Genomics # 1000285) following manufacturer recommendations. Nuclei were loaded onto a Next GEM Chip J (10X Genomics # 1000230), targeting a yield of 3,000 – 6000 nuclei. Sequencing reads were demultiplexed and aligned to the hg38 reference genome using the 10X Genomics CellRanger software (arc v2.0.0) mkfastq function with default settings, and counts were generated using the CellRanger-arc count function. Please see supplemental methods for the bioinformatics approaches to data analysis and interpretation.

#### DNA methylation

5 μm tissue sections were cut from formalin-fixed paraffin-embedded surgically derived specimens from Cushing’s disease adenomas. Autopsy-derived human pituitary gland sections were used as controls as previously published ^28^. DNA was isolated and purified (Qiagen DNeasy #69556). After sodium bisulfite conversion using the Zymogen EZ DNA Methylation kit (#D5001), 1ug each of methylated and unmethylated DNA was analyzed using the Illumina Infinium MethylationEPIC chip (#20087706). Data was processed and analyzed using the MissMethyl R package. ^90^For the *PPP1R17* promoter, 2uL of eluted DNA was used for each PCR reaction, with methylated and unmethylated primers. qRT-PCR was performed using the Sso Universal IT SYBR green supermix (BioRad #1725271). Beta values were quantified using the ratio of methylated / (methylated + unmethylated) expression.

#### Bulk RNAseq

Total RNA was extracted from stable cell lines using the RNeasy Mini Kit (Qiagen, #74004). PolyA+ mRNA isolation, size fragmentation, cDNA synthesis, size selection, and next gen sequencing were performed at the National Intramural Sequencing Center, Bethesda, MD according to their standard protocols, as described previously ^28^. The Illumina TruSeq RNA Sample Prep V2 Kit was used according to manufacturer’s instructions. Analysis of bulk RNA sequencing data was performed using the EdgeR package on total exon counts per gene. A paired analysis was performed with GLM-based approach with design matrix including sample type.

#### Reverse chromatin immunoprecipitation (rChIP)

rChIP was performed by modifying a previously published protocol ^91^. Negative controls included the dCas9-3XFLAG-Biotin Protein, however the synthetic guide RNAs for promoter-specific targeting were excluded. Purification and reversal of cross linking was performed using an EZ-Magna ChIP kit (Millipore Sigma #17-10086). To validate locus-specific pulldown, an aliquot of the eluent was treated with proteinase K (Millipore #20-298) and PCR was run using primers specific to the *PPP1R17* promoter. Protein fragments were then purified and treated with RNAse A (Millipore #20-297) and quantified using Mass spectrometry (SimulTOF 300).

#### Proteomic and phospho-proteomic profiling

LC-MS/MS data acquisition was performed on a Thermo Scientific Orbitrap Lumos mass spectrometer which was coupled to a Thermo Scientific Ultimate 3000 HPLC. Pairwise analysis of total proteins and phosphopeptide/total protein ratios were computed for each tumor-normal pair and compared using two-sample tests of proportions with FDR corrections for multiple testing as appropriate. Phosphoprotein motif analysis was performed using the MEME suite motif-based sequence analysis tool ^92^.

#### Multiplex immunocytochemistry

5 µm-thick paraffin-embedded formalin-fixed human sections were imaged using an Axio Imager.Z2 slide scanning epifluorescence microscope (Zeiss) equipped with a 20X/0.8 Plan-Apochromat (Phase-2) non-immersion objective (Zeiss). Pseudocolored stitched images were exported to Adobe Photoshop and overlaid as individual layers to create multicolored merged composites.

#### Immunohistochemistry

Paraffin embedded tissue blocks were sectioned (5 μm) and stained with an automated immunostainer (Bond-Max, Leica). Sections were imaged on a Nikon Eclipse Ni microscope (Nikon) equipped with a Retiga EXi Fast1394 camera (QImaging) or with a Nikon Eclipse TE200 spinning disk microscope (Nikon) equipped with an EMCCD ImageM digital camera (Hamamatsu), using the iVision software (BioVision).

#### Spatial quantification of mIHC co-expression

Nuclear segmentation of multiplex immunohistochemistry TIFF files was accomplished using ‘Cell Detection’ in Qupath on the DAPI channel. Exported single-cell measurements were imported into Python using pandas and structured using anndata. Tumor regions were defined computationally by calculating a per-cell TumorScore as the mean expression of CIP2A, CYCLIN D1, EGFR, HK2, Kip1, LDHA, NOXA, PCSK1, PFKFB3, PME1, POMC, RAB3B, S100B, SET, SOX2, SOX9, TBX19, cMyc, and RBC, excluding PP2AC and PPP1R17. An adaptive intensity threshold was determined using Otsu’s method from sklearn and cells exceeding this threshold were designated as tumor candidates. Data visualization, including threshold histograms and grouped bar plots with significance annotation, was performed using matplotlib and seaborn. Figures were constructed using Adobe Illustrator.

#### RNA extraction and quantitative real time PCR (qRT-PCR)

RNA was extracted from both human and mouse cells using an RNeasy Mini Kit (Qiagen #74004) according to the manufacturer’s instructions. Complementary DNA (cDNA) libraries were constructed using SuperScript III for qRT-PCR (Invitrogen Life Technologies). Quantitative real-time polymerase chain reaction (qRT-PCR) was performed using the Sso Universal IT SYBR green supermix (BioRad #1725271). Relative gene expression was calculated using the DDCt method with beta actin or GAPDH as housekeeping genes.

#### Cell cycle analysis

Cell cycle was analyzed using a Click-iT Plus EdU Alexa Fluor 647 Flow Cytometry assay kit (Thermo Fisher #C10635) following manufacturer’s instructions. Proportions of cells in the S-phase were compared between samples using two-sample tests of proportions, with P < 0.05 considered statistically significant.

#### Cell viability assays

Cell viability assay was performed using CellTiterGlo (Promega #G7571) according to the manufacturer’s instructions. Plates were incubated for 30 minutes at room temperature prior to recording luminescence on a Synergy Neo2 microplate reader (BioTek).

#### Proximity ligation assay (PLA)

Proximity between PPP1R17 and PP2Ac was determined using a Duolink Proximity Ligation Assay (PLA) Red Mouse/Rabbit kit (Sigma #DUO92101) following the manufacturer’s instructions. 5 μm human CD adenoma sections were processed and mounted using Duolink in-situ mounting media with DAPI and imaged.

#### PP2A activity assay

Human PPP1R17 and PP2Ac were cloned into pET FLAG vectors, transformed using BL21 gold competent bacterial cells (Sigma) and amplified. After amplification, FLAG-tagged PPP1R17 and PP2Ac proteins were purified using a HiTrap nickel column (Sigma). PP2A activity was assessed using a Serine/Threonine Phosphatase Assay (Promega #V2460) following manufacturer’s instructions. Plates were incubated for 30 minutes at room temperature prior to quantifying absorbance of the molybdate:malachite green:phosphate complex at 630nm on a Synergy Neo2 microplate reader (BioTek).

#### Thermal shift assay

Protein thermal shift assays were performed on 384-well plate or a QantaBio qPCR instrument. The plate was subjected to thermal shift by ramping the temperature from 25°C to 99°C at 0.05°C increments per second. The relative fluorescence emitted by the thermal shift dye was recorded during the temperature ramp phase and plotted versus temperature. The Tm values were plotted as a function of TDP43 concentration in GraphPad Prism, and the inflexion point of the curve determined the binding affinity of ligand to binding protein. The experiments were completed with n = 4 replicate samples per treatment. Each experiment was repeated 3 times.

#### AlphaFold multimer

AlphaFold multimer was designed specifically to predict the structures of protein complexes more accurately than AlphaFold and AlphaFold2 ^53^. We generated protein structure and interaction inference using the AlphaFold2 algorithm ^93^. Briefly, FASTA files with the canonical protein sequence (PP2A, PPP1R17, and PME1) were analyzed on the NIH HPC cluster (BioWulf). The AlphaFold2 algorithm preferentially used existing experimental PDB structures where available. Multiple sequence alignment and model predictions were then generated for protein interaction inference and multimer models. Multimer model prediction alignment error rates, and pLDDT values were further analyzed with the alphapickle package (M. J. Arnold. 2021. AlphaPickle. doi.org/10.5281/zenodo.5708709). The top ranked multimer models were used to analyze protein interaction sites, and to generate the figure panels.

#### Statistical analysis

Means were compared using two-sample T tests after testing for variance equality and using Welch’s approximation for degrees of freedom ^94^. P values < 0.05 two-tailed were considered statistically significant. Statistical analyses were performed using STATA 14/IC (StataCorp LP, College Station, Texas) or GraphPad Prism 9.0 (GraphPad Software, La Jolla, California).

