## Supplemental Figures for "Phosphoproteomic dysregulation drives tumor proliferation in Cushing’s disease"

### Canonical cell types identified in each sample

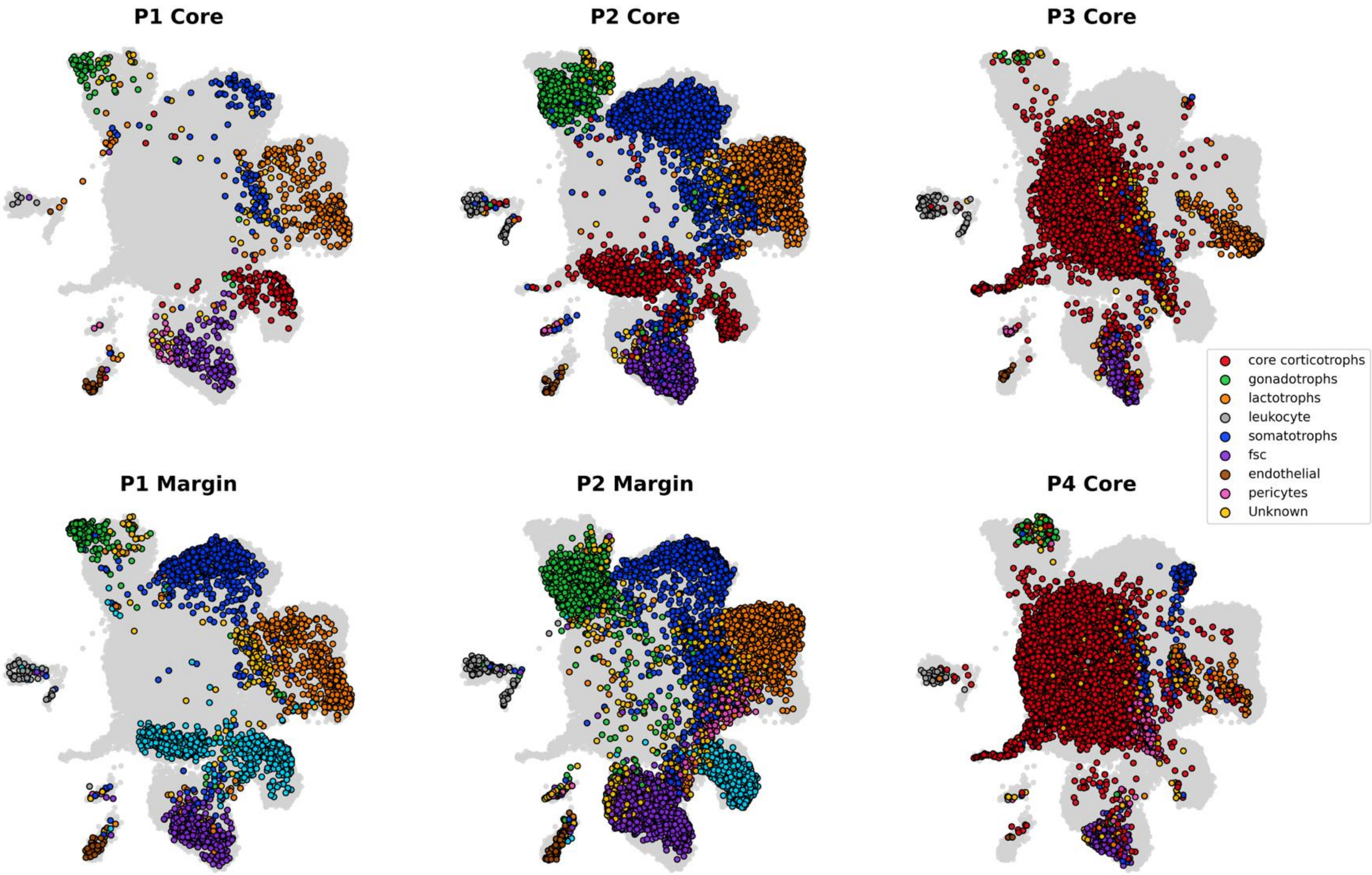


**Supplemental Figure S1.** UMAPs of canonical cell types in each individual tumor sample. Refer to Figure 1D for aggregate figure.

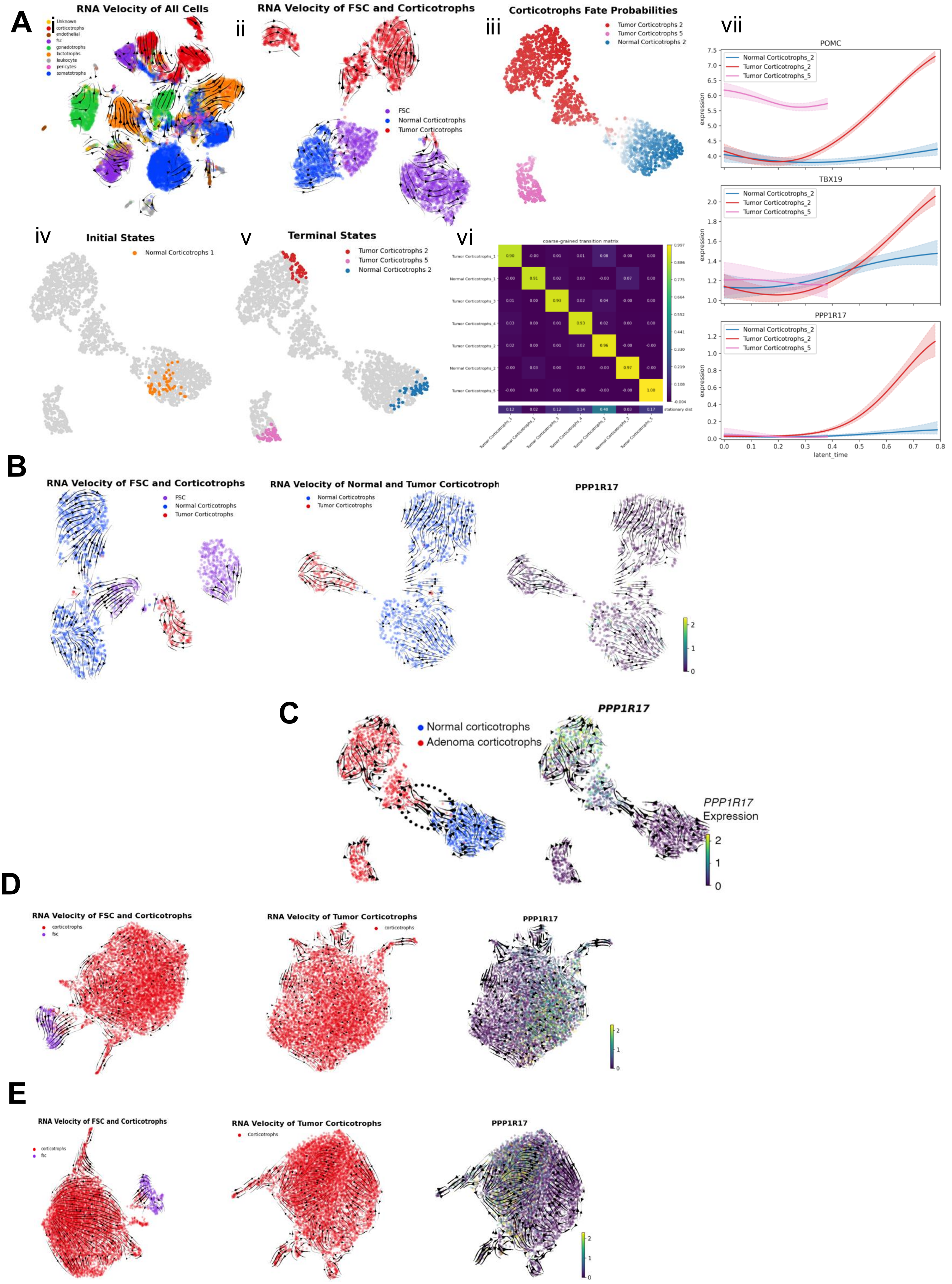

A
i
RNA Velocity of All Cells
Unknown
corticotrophs
endothelial
fsc
gonadotrophs
lactotrophs
leukocyte
pericytes
somatotrophs
ii
RNA Velocity of FSC and Corticotrophs
FSC
Normal Corticotrophs
Tumor Corticotrophs
iii
Corticotrophs Fate Probabilities
Tumor Corticotrophs 2
Tumor Corticotrophs 5
Normal Corticotrophs 2
vii
POMC
TBX19
PPP1R17
Normal Corticotrophs_2
Tumor Corticotrophs_2
Tumor Corticotrophs_5
expression
latent_time
iv
Initial States
Normal Corticotrophs 1
v
Terminal States
Tumor Corticotrophs 2
Tumor Corticotrophs 5
Normal Corticotrophs 2
vi
coarse-grained transition matrix
stationary dist
B
RNA Velocity of FSC and Corticotrophs
FSC
Normal Corticotrophs
Tumor Corticotrophs
RNA Velocity of Normal and Tumor Corticotroph
Normal Corticotrophs
Tumor Corticotrophs
PPP1R17
C
Normal corticotrophs
Adenoma corticotrophs
PPP1R17
PPP1R17
Expression
D
RNA Velocity of FSC and Corticotrophs
corticotrophs
fsc
RNA Velocity of Tumor Corticotrophs
corticotrophs
PPP1R17
E
RNA Velocity of FSC and Corticotrophs
corticotrophs
fsc
RNA Velocity of Tumor Corticotrophs
Corticotrophs
PPP1R17

**Supplemental Figure S2.** Tumor corticotroph lineage tracing analysis. **A.** CellRank and scVelo analysis of patient sample P2. **i.** scVelo velocities projected onto UMAP embedding of all cells in both tumor and normal compartments colored by cell type. **ii.** scVelo velocities projected onto UMAP embedding of tumor corticotrophs, normal corticotrophs, and folliculostellate cells (FSC). **iii.** UMAP of corticotrophs from tumor and normal compartments. Cells are colored by CellRank fate probabilities, which is the likelihood that a given cell will transition toward the terminal population. **iv.** CellRank identified initial cell states. **v.** CellRank identified terminal cell states. **vi.** Transition matrix showing the fate probabilities of each macrostate. **vii.** Relative expression of relevant genes in each macrostate plotted over latent time. **B.** CellRank and scVelo analysis of patient sample P1. scVelo velocities projected onto UMAP embedding of tumor corticotrophs, normal corticotrophs, and FSCs (left), scVelo velocities projected onto UMAP embedding of tumor and normal corticotrophs (center), scVelo velocities projected onto UMAP embedding of tumor and normal corticotrophs with cells colored by relative *PPP1R17* expression (center). **C.** CellRank and scVelo analysis of patient sample P2. scVelo velocities projected onto UMAP embedding of tumor and normal corticotrophs. Cells colored by relative *PPP1R17* expression. **D.** CellRank and scVelo analysis of patient sample P3. scVelo velocities projected onto UMAP embedding of tumor corticotrophs and FSCs (left), scVelo velocities projected onto UMAP embedding of only corticotrophs (center), scVelo velocities projected onto UMAP embedding of corticotrophs with cells colored by relative *PPP1R17* expression (center). **E.** CellRank and scVelo analysis of patient sample P3. scVelo velocities projected onto UMAP embedding of tumor corticotrophs and FSCs (left), scVelo velocities projected onto UMAP embedding of only corticotrophs (center), scVelo velocities projected onto UMAP embedding of corticotrophs with cells colored by relative *PPP1R17* expression (center).

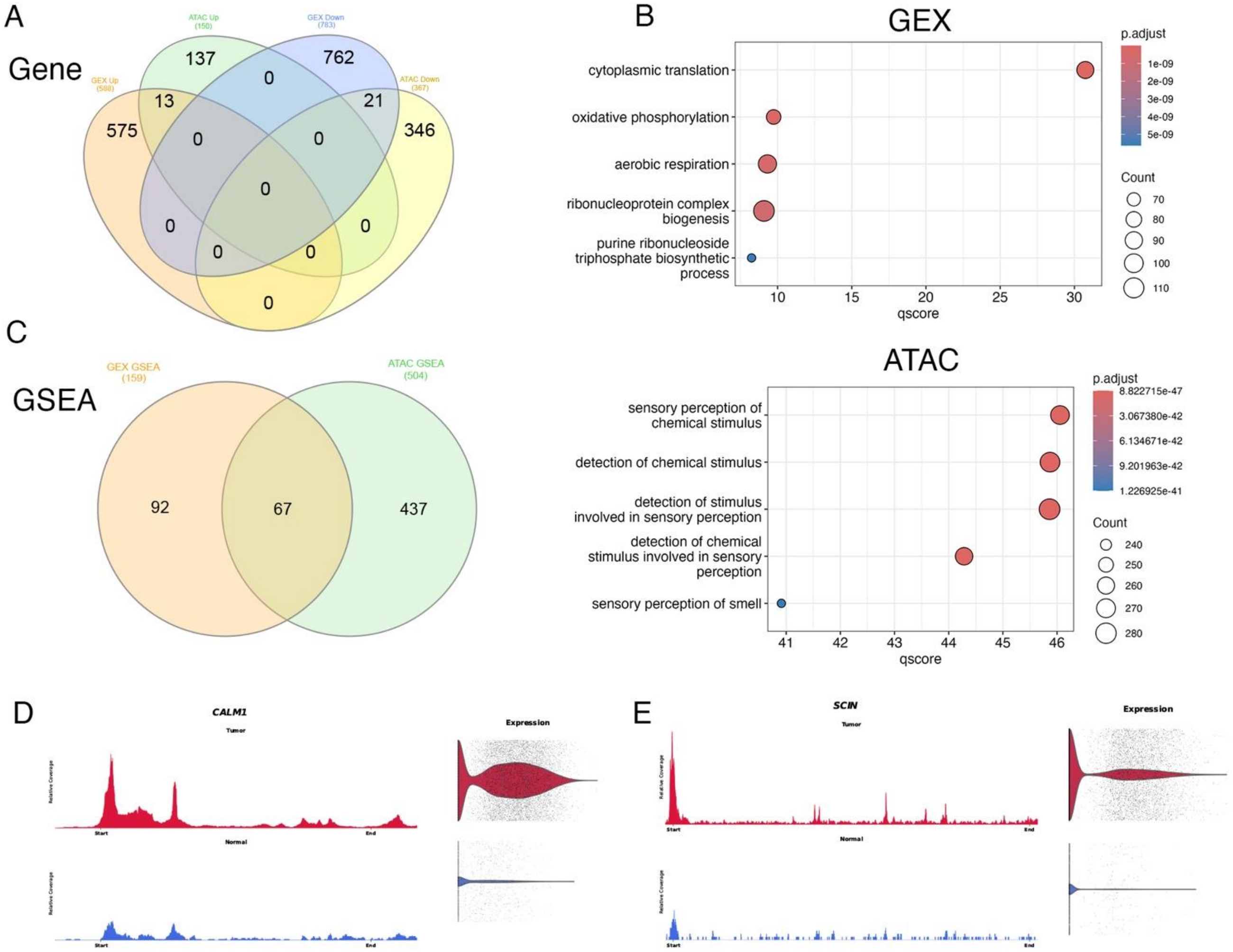


**Supplemental Figure S3**. **A.** Multiome analysis comparing snRNAseq and snATACseq in CD corticotrophs versus adjacent normal corticotrophs. **B.** GSEA analysis of pathways upregulated by snRNAseq in CD corticotrophs versus adjacent normal corticotrophs. **C.** Left panel: overlap between genes differentially expressed (GEX) and genes differentially accessible (ATAC) in CD corticotrophs. Right panel: GSEA analysis of pathways differentially accessible by snATACseq in CD corticotrophs. **D.** Comparison of chromatin accessibility (left panels) and gene expression (right panels) for NF-Y target gene CALM1 in CD corticotrophs (red, tumor) versus adjacent normal corticotrophs (blue, normal). **E.** Differential chromatin accessibility (left panels) and gene expression (right panels) for SP1 target gene SCIN in CD corticotrophs (red, tumor) versus adjacent normal corticotrophs (blue, normal).

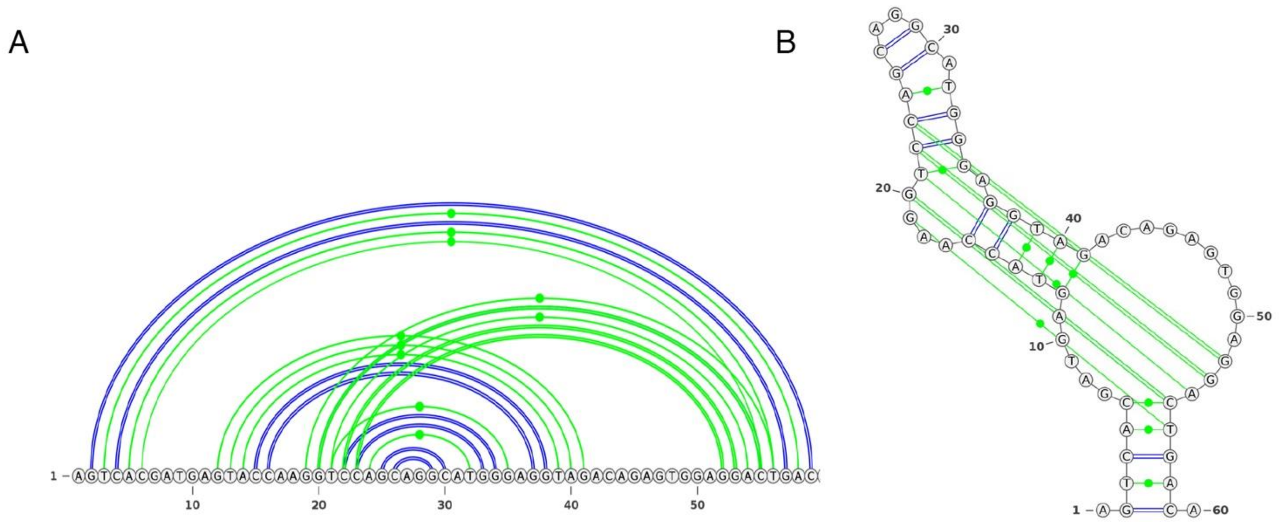


**Supplemental Figure S4**. The novel lncRNA PPP1R17-203 is transcribed from an internal *PPP1R17* site. **A.** Long-range interactions predicted for PPP1R17-203. **B.** Predicted 3D structure of PPP1R17-203. Source: https://bioinformaticslab.erc.monash.edu/linc2function

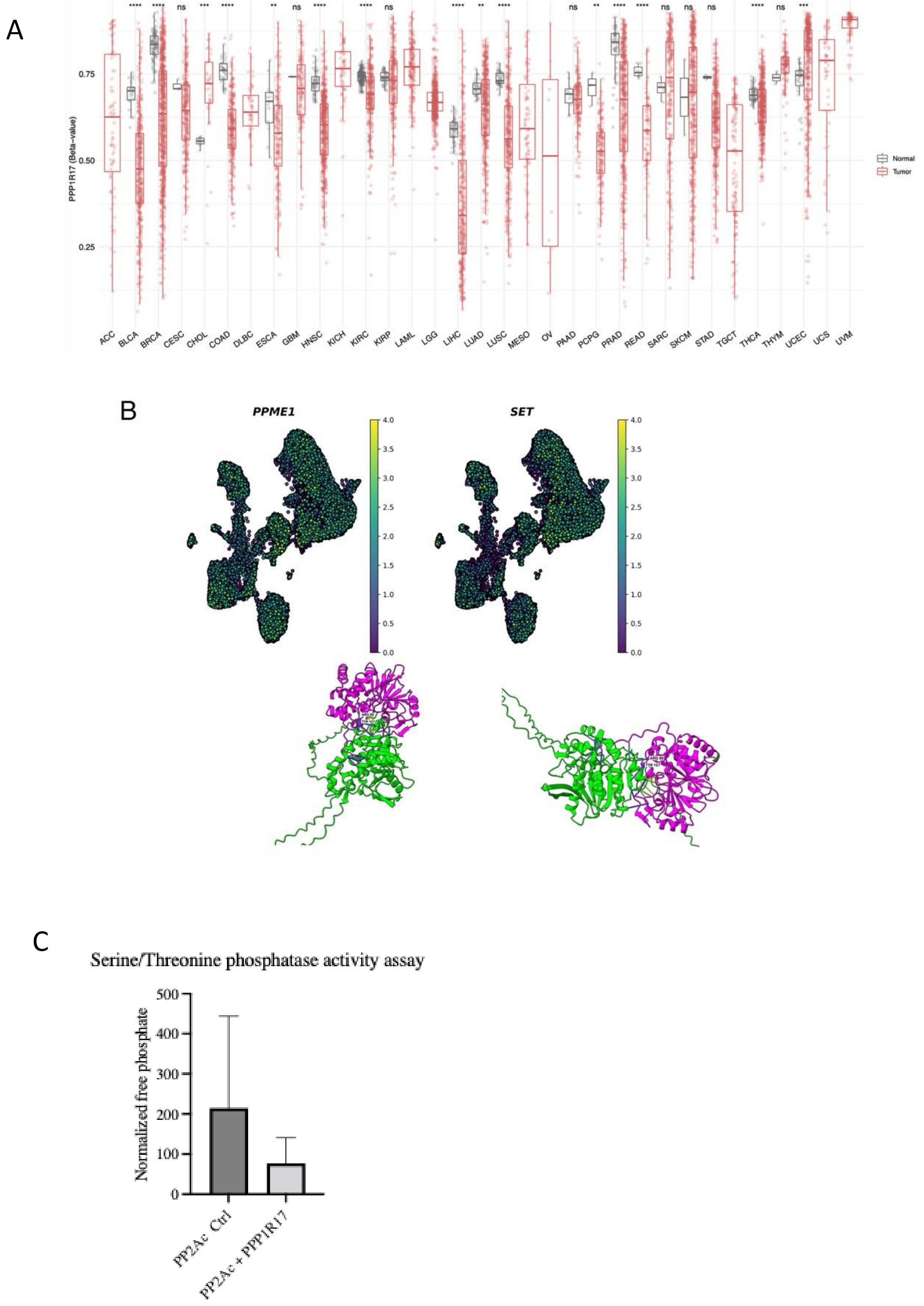


**Supplemental Figure S5. A.** DNA methylation analysis from the TCGA database demonstrating *PPP1R17* promoter hypomethylation in multiple solid tumors including hepatocellular carcinoma and head and neck squamous cancer. **B.** Endogenous PP2A agonists*PPME1* and *SET* are not transcriptionally overexpressed in CD. UMAP projections demonstrating expression patterns of *PPME1* and *SET* in *POMC*-overexpressing CD adenoma cells. AlphaFold multimer predicts PME1 and SET interaction with PP2Ac at Arg89 and share an additional interaction site at Tyr127 with PPP1R17. **C.** Serine/threonine phosphatase assay in cell-free media comparing phosphatase activity of purified PP2Ac in the presence or absence of PPP1R17. UMAP = uniform manifold approximation and projection. CD = Cushing's disease.

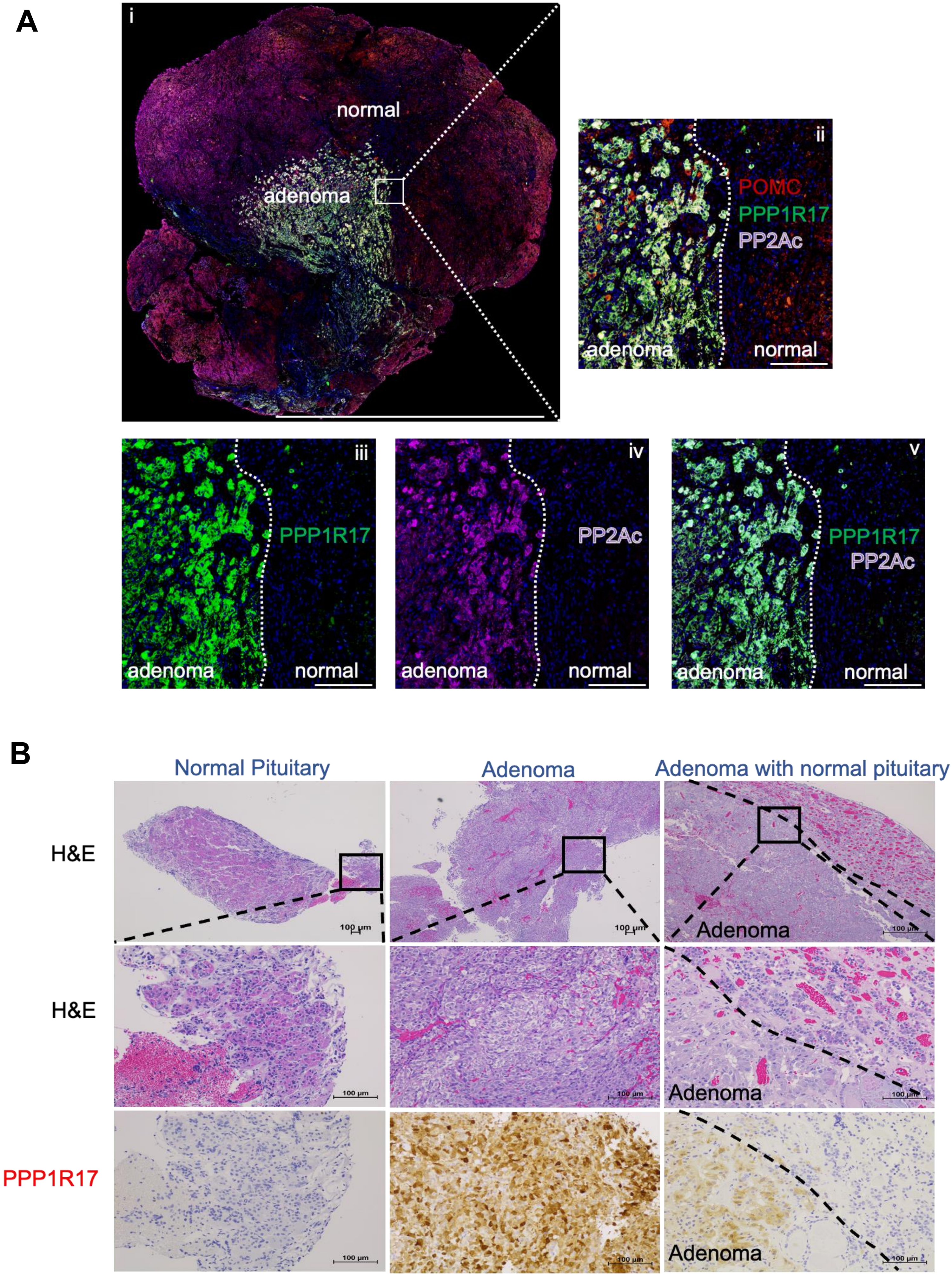
A
i
normal
adenoma
ii
POMC
PPP1R17
PP2Ac
adenoma
normal
iii
PPP1R17
adenoma
normal
iv
PP2Ac
adenoma
normal
v
PPP1R17
PP2Ac
adenoma
normal
B
Normal Pituitary
Adenoma
Adenoma with normal pituitary
H&E
H&E
PPP1R17
100 μm
Adenoma
Adenoma
Adenoma

**Supplemental Figure S6.** PPP1R17 expression is confined to CD adenomas. **A.** Multiplex immunohistochemistry of CD adenoma and surrounding normal pituitary gland (i) showing PPP1R17 expression (green) compared to corticotroph-specific POMC expression (red). PP2Ac expression (purple) staining is compared to PPP1R17 staining within the adenoma (iii – v). Scale bar = 50µM. **B.** Representative H&E of normal pituitary gland and CD adenoma demonstrating PPP1R17 staining. CD = Cushing's disease. H&E = hematoxylin and eosin.

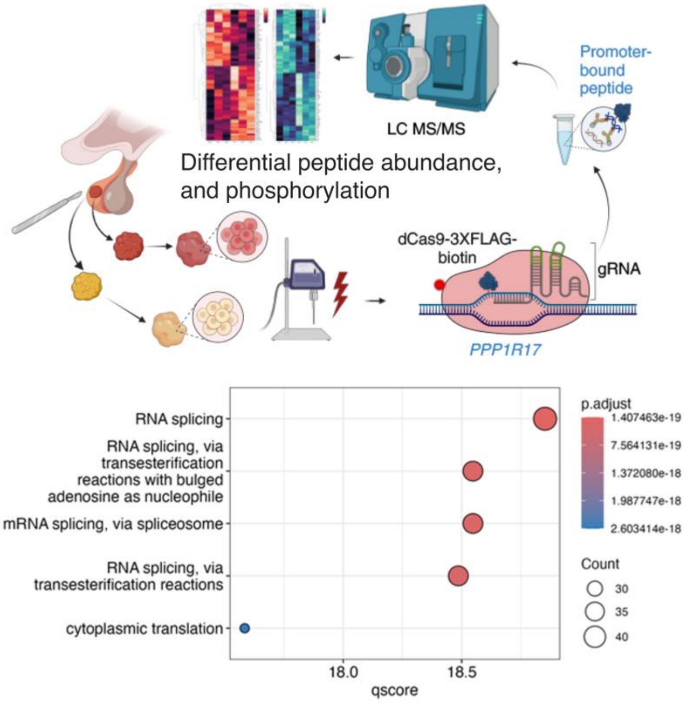


**Supplemental Figure S7.** Schematic for reverse-ChIP performed using CRISPR dCas9-3XFLAG-biotin and a gRNA targeting the *PPP1R17* promoter to pull down promoter-bound peptides for mass spectrometry. GSEA analysis for proteins bound to the *PPP1R17* promoter reveals enrichment for RNA splicing and RNA processing.

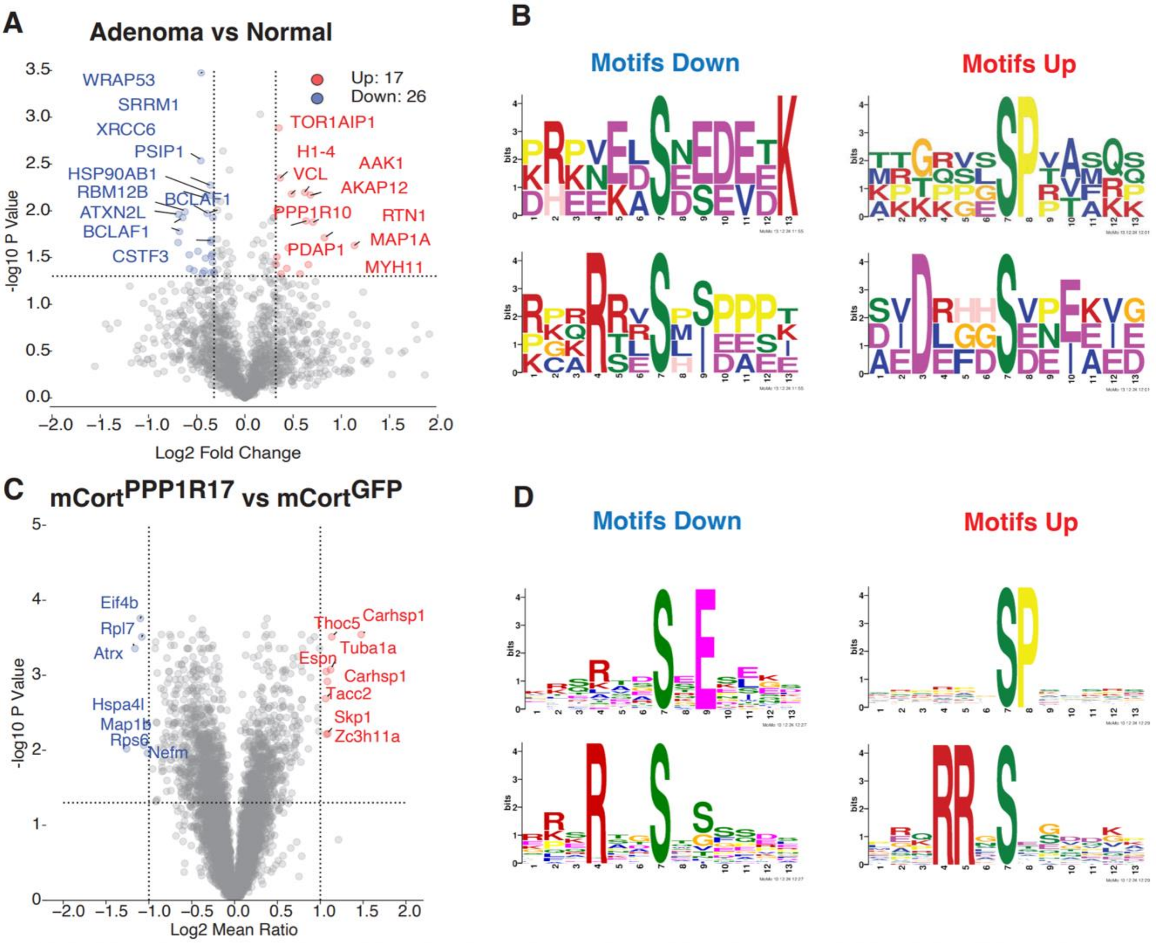


**Supplemental Figure S8**. **A.** TMT LC-MS/MS quantification of phosphopeptide/peptide ratios in core CD adenomas versus margin (n = 3/group) shows several hyperphosphorylated and dephosphorylated targets. **B**. MoMo motif analysis of preferentially hyperphosphorylated and dephosphorylated motifs in CD adenoma core versus margin (n = 3/group). **C**. TMT LC-MS/MS quantification of phosphopeptide/peptide ratios in mCortPPP1R17 versus mCortGFP cells (n = 5/group). **D.** MoMo motif analysis of preferentially hyperphosphorylated and dephosphorylated motifs in mCortPPP1R17 versus mCortGFP cells (n = 5/group).

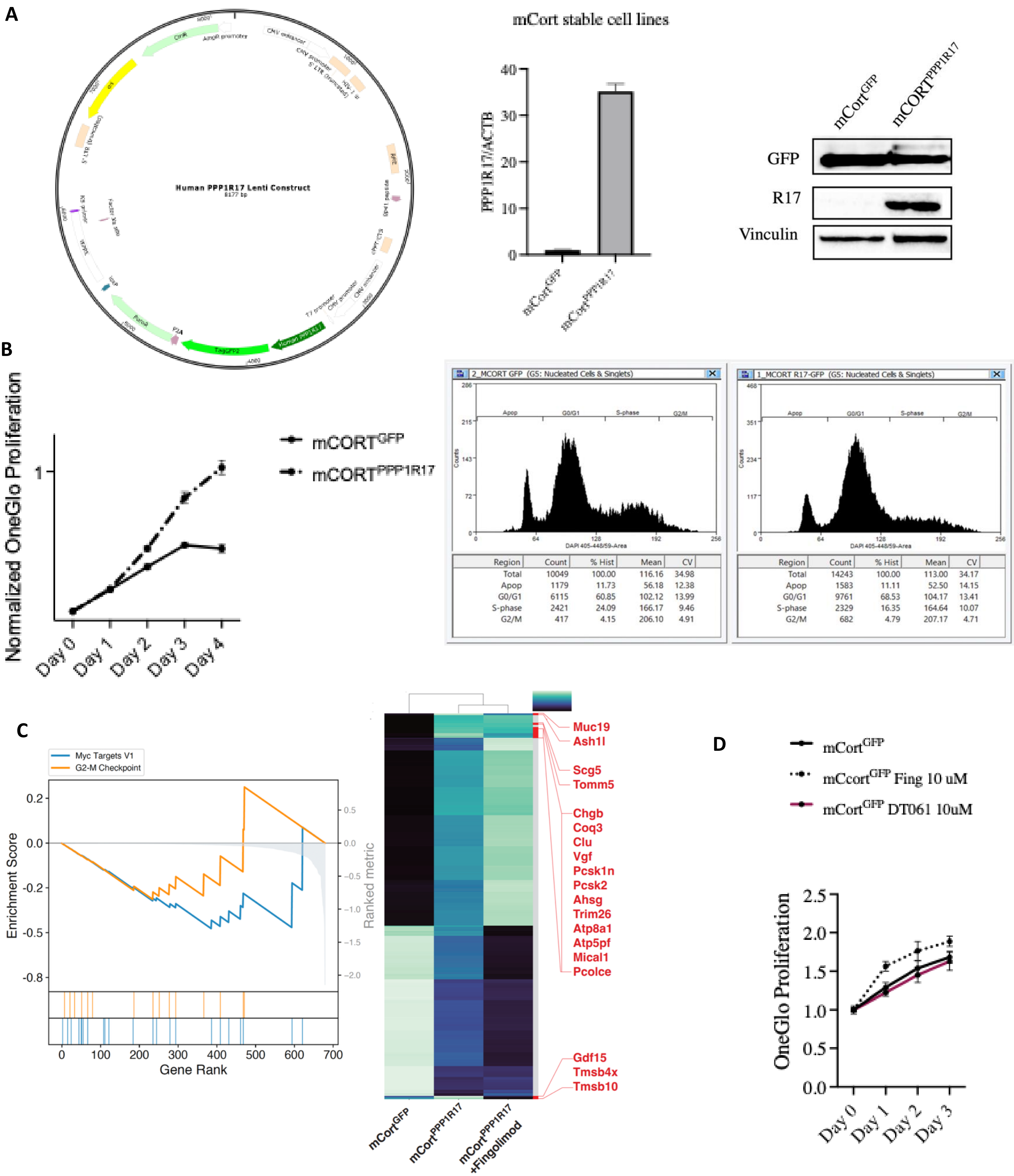


**Supplemental Figure S9**. **A**. Lentiviral vector overexpressing *PPP1R17* tagged to *GFP*. Stable transfection in mCort cells was verified by qRT-PCR and by Western immunoblotting. **B.** Stable *PPP1R17* overexpression in mCort cells (mCort$^{PPP1R17}$) showed hyperproliferation compared to mCort$^{GFP}$ cells (n = 8/group). mCortPPP1R17 cells also showed acceleration of the cell cycle. **C.** GSEA analysis of protein pathways and heatmap of individual proteins hyperphosphorylated in mCort$^{PPP1R17}$ but rescued by 48-hour fingolimod treatment. Results of TMT labeling and mass spectrometry; n = 5/group; one-way ANOVA $P < 0.05$). **D.** Effect of fingolimod and DT061 on proliferation in mCort$^{GFP}$ cells; n = 8/group.

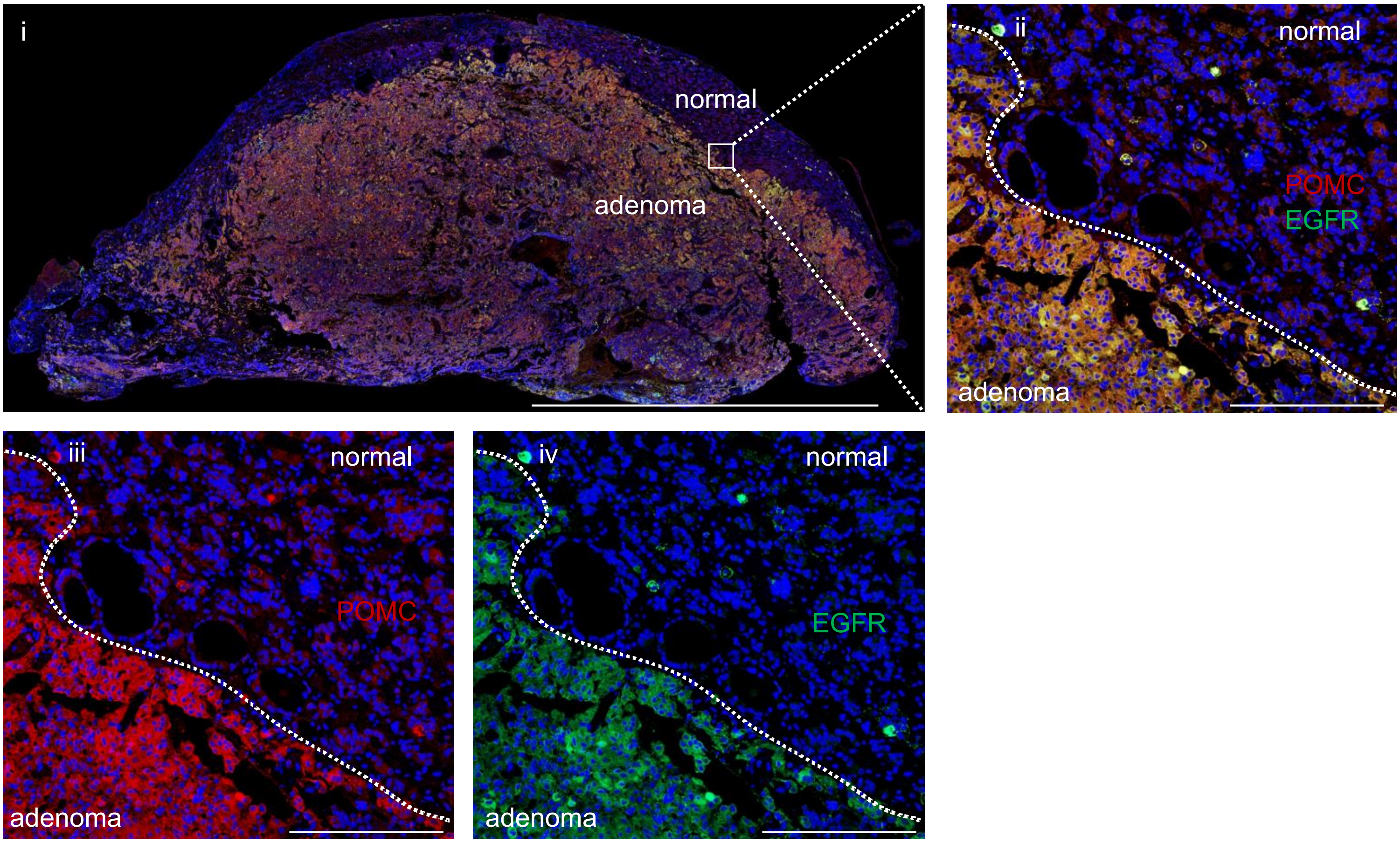


**Supplemental Figure S10**. EGFR is overexpressed in CD. Representative mIHC image of whole-mounted CD adenoma and adjacent normal gland (i) with corticotroph-specific POMC expression. (ii-iv) EGFR expression (green) was compared to POMC expression (red). Scale bar = 200 μM.

### Supplemental Figure S9

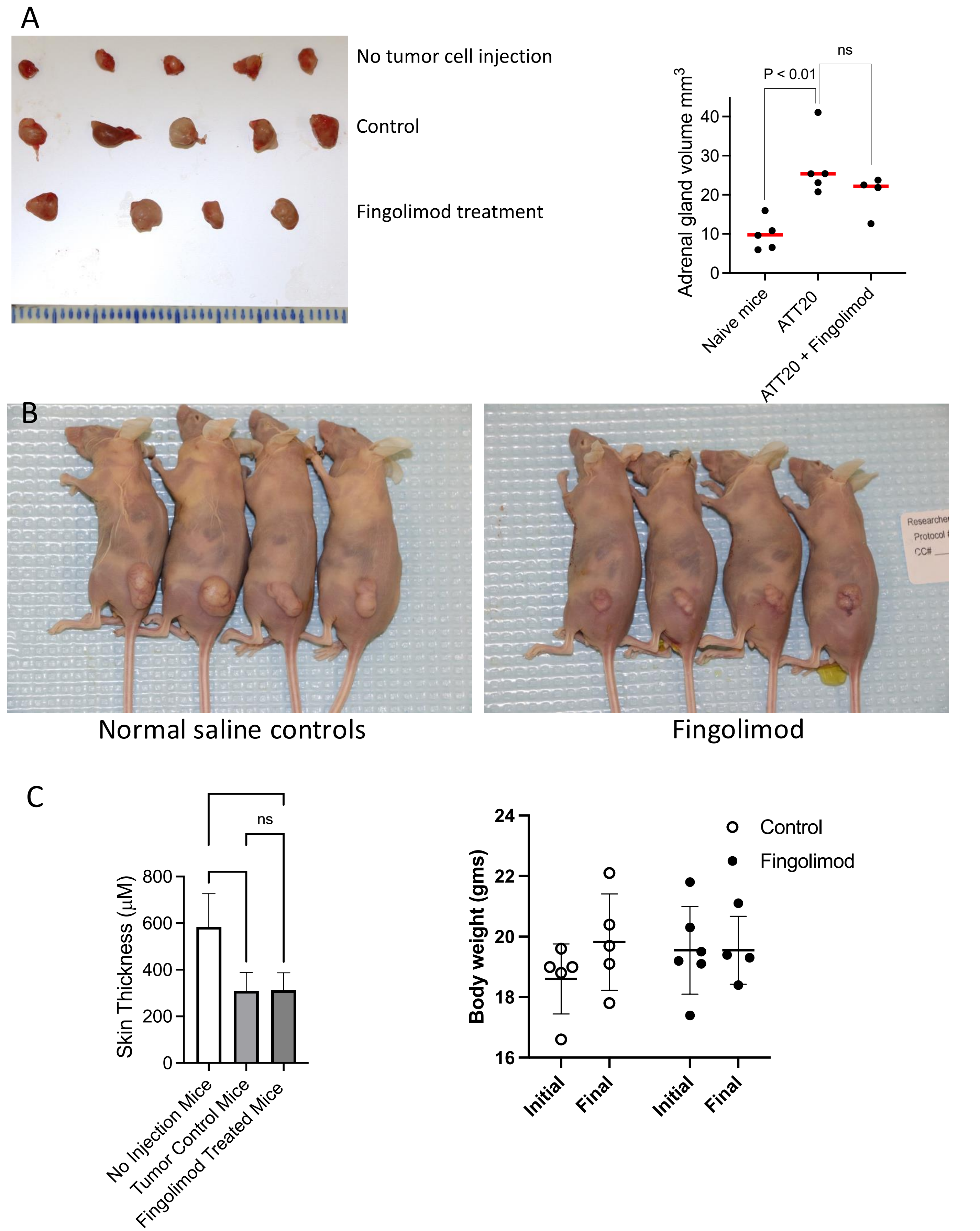


**Supplemental Figure S11**. **A**. Implantation of ATT20 cells into the flanks of nude mice induced adrenal gland hypertrophy (n = 5/group), which did not return to baseline after fingolimod treatment (n = 4). **B.** Nude mice bearing ATT20 flank xenografts formed tumors after 10-12 days. After tumor establishment, intraperitoneal fingolimod 1mg/kg daily for 17 days led to decreased tumor size. **C.** Fingolimod treatment had no effect on skin thickness or animal body weight.
